# Native Elongation Transcript sequencing reveals temperature dependent dynamics of nascent RNAPII transcription in *Arabidopsis*

**DOI:** 10.1101/834507

**Authors:** Peter Kindgren, Maxim Ivanov, Sebastian Marquardt

**Author notes:** indicates equal contribution (Peter Kindgren, Maxim Ivanov).

## Abstract

Temperature profoundly affects the kinetics of biochemical reactions, yet how large molecular complexes such as the transcription machinery accommodate changing temperatures to maintain cellular function is poorly understood. Here, we developed plant native elongating transcripts sequencing (plaNET-seq) to profile genome-wide nascent RNA polymerase II (RNAPII) transcription during the cold-response of *Arabidopsis thaliana* with single-nucleotide resolution. Combined with temporal resolution, these data revealed transient genome-wide reprogramming of nascent RNAPII transcription during cold, including characteristics of RNAPII elongation and thousands of non-coding transcripts connected to gene expression. Our results suggest a role for promoter-proximal RNAPII stalling in predisposing genes for transcriptional activation during plant-environment interactions. At gene 3’-ends, cold initially facilitated transcriptional termination by limiting the distance of read-through transcription. Within gene bodies, cold reduced the kinetics of co-transcriptional splicing leading to increased intragenic stalling. Our data resolved multiple distinct mechanisms by which temperature transiently altered the dynamics of nascent RNAPII transcription and associated RNA processing, illustrating potential biotechnological solutions and future focus areas to promote food security in the context of a changing climate.

## INTRODUCTION

Changes to ambient temperatures challenge the development and growth of living organisms. While mammals retain a stable body temperature, sessile organisms such as plants continually sense their environment and rely on molecular mechanisms that compensate for temperature changes(1). Alterations to the ambient temperature frequently lead to re-programming of the transcriptional output by RNA polymerase II (RNAPII) that reflects steady-state levels of messenger RNAs and non-coding RNAs in the cell(2,3). Sequence-specific transcription factors controlling the initiation of transcription often shape these responses. However, the significance of mechanisms regulating eukaryotic gene expression after initiation, for example through control of elongation of the nascent RNA chain is increasingly appreciated(4). Genome-wide profiling of transcriptionally engaged RNAPII complexes has identified low-velocity regions of RNAPII elongation at the beginning (i.e. promoter-proximal stalling) and the end (i.e. poly-(A) associated stalling) of genes(4,5). Organisms appear to alter the activity of RNAPII at these regions to re-program their transcriptional output to acclimate to temperature changes. The release from promoter-proximal stalling at heat-shock genes facilitates rapid transcriptional induction in response to heat in *Drosophila*(6), and promoter-proximal stalling is reduced genome-wide when temperatures increase in mammalian cell cultures(7). RNAPII accumulation at gene ends is associated with the mechanism of transcriptional termination(8). Here, molecular complexes associated with nascent RNAPII transcript cleavage at the poly(A)-signal (PAS) regulate RNAPII activity to ensure accurate processing of the nascent transcript(8). RNAPII continues to transcribe past the PAS until 5’-to-3’ exonucleases catch up with transcribing RNAPII to mediate transcriptional termination(8–10). Hence, transcriptional termination is determined by kinetic competition between the speed of RNAPII transcription after nascent transcript cleavage and the termination factor(11). Temperature increases lengthen the read-through transcription distance at gene ends in several organisms(11,12), suggesting connections between temperature, RNAPII stalling at gene borders and the efficiency of transcriptional termination. However, the immediate genome-wide effects of low temperatures on nascent RNAPII transcription in eukaryotes are unclear.

Transcriptionally engaged RNAPII complexes can be visualized by Native Elongating Transcript sequencing (NET-seq)(13–16). NET-seq provides a strand-specific snapshot of nascent RNAPII transcription at single-nucleotide resolution genome-wide(16). The capture of nascent RNA by NET-seq enables the detection of RNAs that are usually subjected to co-transcriptional RNA degradation. This advantage of NET-seq helps to detect long non-coding RNAs (lncRNAs), as these tend to be targeted for co-transcriptional RNA degradation by the nuclear exosome RNA degradation complex(17,18). Moreover, NET-seq in yeast and mammals allowed estimates of the average length of cryptic read-through transcription that allows quantitative analyses of the transcription termination mechanism(19,20). An additional advantage of NET-seq data are insights into co-transcriptional RNA splicing, since part of the spliceosome is co-purified with transcribing RNAPII complexes(15,21). Nascent RNAPII transcription slows down close to exon-intron boundaries in a splicing-dependent manner and is responsible for intragenic RNAPII stalling(15). Splicing regulation is essential for the cold-response in *Arabidopsis*(22,23) but how this is connected to molecular adjustments of nascent RNAPII transcription is largely unknown.

Here, we developed a NET-seq approach to study nascent transcription in the model plant *Arabidopsis thaliana* (plaNET-seq). We analyzed the temporal dynamics of nascent RNAPII transcription in response to cold. Our data revealed transient molecular adaptations of transcription that include changes to promoter-proximal stalling, elongation, termination and many novel non-coding transcription events overlapping gene expression domains. Our data provide genome-wide support for a transient re-programming of nascent RNAPII transcription during cold exposure, highlighting a cellular compensation mechanism at the level of nascent RNAPII transcription to assist optimal growth of multicellular organisms in challenging environments.

## MATERIALS & METHODS

### Plant material and growth conditions

*A. thaliana* seeds were surface-sterilized in ethanol and grown on ½ MS + 1% sucrose media in long day conditions (16 h light/8 h dark) at 22°C/18°C. Light intensity during day hours was approximately 100 μE m^−2^ s^−1^. 10-day old seedlings were used for all experiments. The NRPB2-FLAG line was described in(24). The construct covers a lethal *nrpb2-1* allele (SAIL_859B04). The *fas2-4* mutant is described in(25). For inhibition of splicing, seedlings were grown on filter paper covered ½ MS + 1% sucrose for 10 days then transferred to DMSO, 5 μM pladienolide B (Santa Cruz) or 5 μM Herboxidiene (Focus Biomolecules) containing plates for 6 or 24 hours. For low temperature treatment, 10-day old seedlings were transferred to 4°C and approximately 25 μE m^−2^ s^−1^ for indicated times.

### Total RNA isolation and RT-qPCR

Total RNA was isolated from Arabidopsis seedlings grown for 10 days and exposed to DMSO or splicing inhibitors for 6 or 24 hours with RNeasy Plant Mini Kit (Qiagen) according to manufacturers’ instructions. 5 μg of total RNA was treated with Turbo DNaseI (Ambion) to remove any genomic DNA. Subsequently, 1 μg of DNase-treated RNA was converted to cDNA using SuperScript IV (Invitrogen) with random primers according to manufacturers’ instructions. Quantitative PCR was performed in 3 technical replicates with the GoTaq qPCR Master Mix (Promega) in 384 well plates. The PCR was run in a CFX384 Touch Real-Time PCR Detection System (BioRad) and monitored by the CFX Manager software (BioRad). Threshold values were subsequently exported to Excel and processed further. All oligos used for the PCR can be found in Supplementary Dataset 3.

### Isolation of nascent RNA

3 grams of seedlings were flash frozen in liquid nitrogen and grinded to fine powder in a mortar. The powder was transferred to a falcon tube with 15 ml NUC1 buffer (0.4 M sucrose, 10 mM Tris-HCl pH 8.0, 10 mM MgCl_2_, 5 mM β-mercaptoethanol, proteinase inhibitor tablet (Roche) and RNase inhibitor (Molox)) and allowed to thaw at 4°C with rotation. After centrifugation (5000 g, 20 min, 4°C), the pellet was dissolved in 1 ml NUC2 buffer (0.25 M sucrose, 10 mM Tris-HCl pH 8.0, 10 mM MgCl_2_, 5 mM β-mercaptoethanol, proteinase inhibitor tablet, RNase inhibitor and 0.3% Tween-20) and centrifuged again (12000 g, 10 min, 4°). The resulting pellet was dissolved in 0.3 ml NUC3 buffer (1.7 M sucrose, 10 mM Tris-HCl pH 8.0, 2 mM MgCl_2_, 5 mM β-mercaptoethanol, proteinase inhibitor tablet, RNase inhibitor and 0.15% Tween-20), placed on top of 0.9 ml clean NUC3 buffer and centrifuged (16000 g, 60 min, 4°C). The purified nuclear fraction was dissolved and lysed in 1.5 ml plaNET-seq lysis buffer (0.3 M NaCl, 20 mM Tris-HCl pH 7.5, 5 mM MgCl_2_, 5 mM DTT, proteinase inhibitor tablet, RNase inhibitor and 0.5% Tween-20). Lysis was performed at 4°C with rotation (2000 rpm), included DNaseI treatment (Invitrogen) and was followed by centrifugation (10000 g, 10 min, 4°C). The supernatant was transferred to a new tube and incubated with Dynabeads M-270 (Invitrogen) bound with anti-FLAG antibody (10 μg, Sigma-Aldrich F3165) for 2 hours at 4°C with gentle rotation. Following 6 times 1 ml washes with wash buffer (0.3 M NaCl, 20 mM Tris-HCl pH 7.5, 5 mM MgCl_2_, 5 mM DTT, proteinase inhibitor tablet and RNase inhibitor), bound proteins were eluted with 3xFLAG peptide (0.5 mg/ml, ApexBio). Elution was performed 2 times with 0.1 ml 3xFLAG peptide for 20 min at 4°C. RNA attached to purified protein complexes was isolated with the miRNeasy kit (Qiagen) according to manufacturer’s instructions. RNA was quantified with RNA Pico kit on Bioanalyzer 2100 (Agilent).

### Preparation of plaNET-seq libraries and sequencing

Libraries were constructed according to Bioo Scientific’s NEXTflex Small RNA-seq kit v3 following a custom protocol. Unlike the original protocol provided by the manufacturer, our custom protocol incorporates RNA fragmentation step in order to avoid underrepresentation of longer molecules of nascent RNA compared to shorter ones (Supplementary fig. 1b). Approximately 100 ng RNA was used for each library. After the ligation of the 3’-linker, RNA was fragmented in alkaline solution (100 mM NaCO_3_ pH 9.2, 2 mM EDTA) to a fragment size of 20-150 bp. After fragmentation, RNA was cleaned up with AMPure RNAclean XP beads, treated with PNK (20 U, NEB) for 20 min at 37°C and then re-annealed with 8 μM RT-primer (70°C, 5 min; 37°C, 30 min; 25°C, 15 min. Oligo sequence: 5’-GCCTTGGCACCCGAGAATTCCA-3’). The RNA was then re-introduced to the manufacturer’s protocol at the adapter inactivation step. For detailed step-by-step library preparation protocol, refer to Supplementary fig. 1b. Depending on the library, 10-16 cycles of PCR was used and the final library was checked with Agilent’s DNA High sensitivity kit on a Bioanalyzer 2100 before sequencing. Libraries were sequenced with the Illumina HiSeq-PE150 platform at Novogene (en.novogene.com).

**Figure 1:**
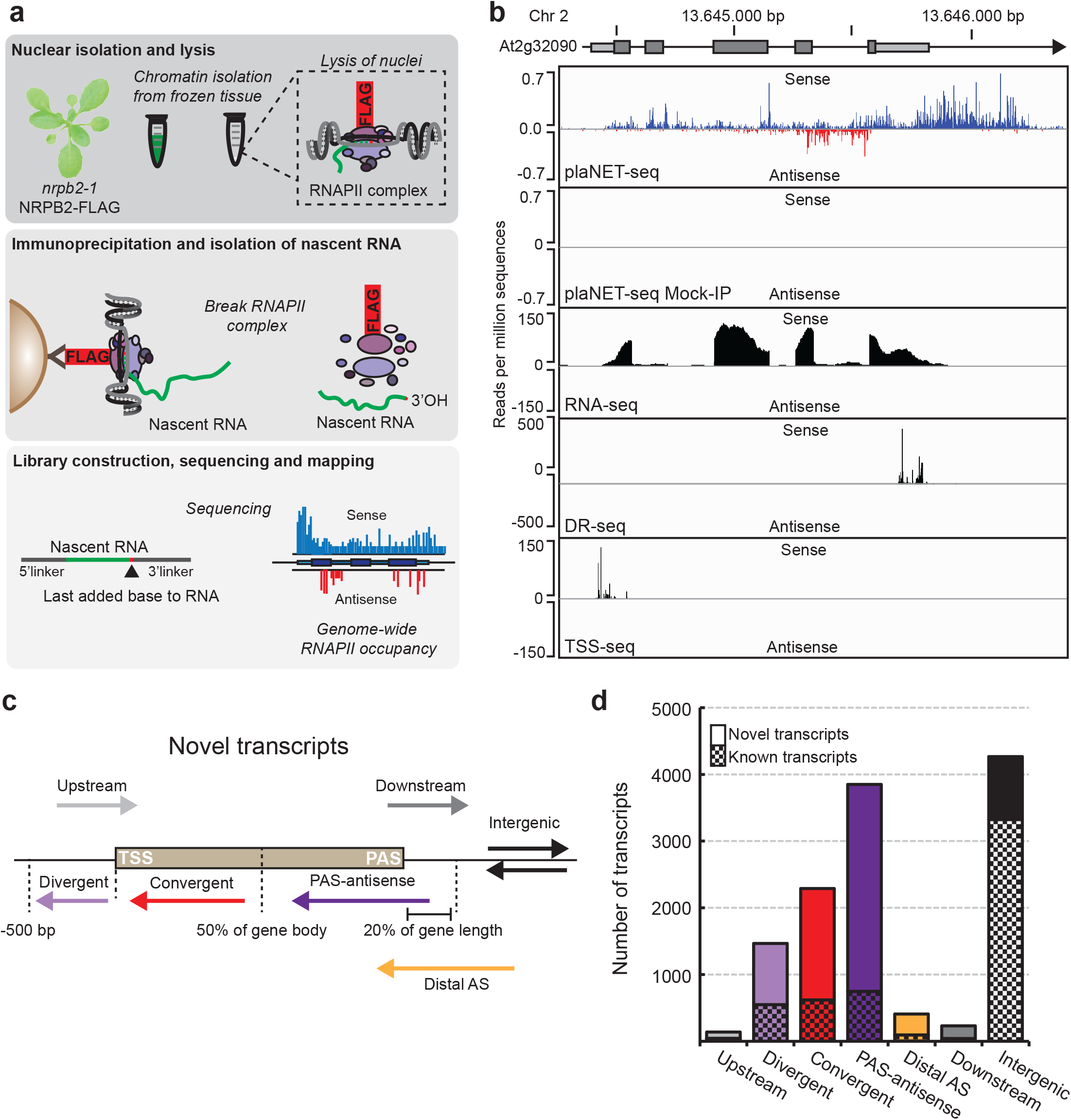
Genome-wide detection of nascent transcription in response to low temperature with plaNET-seq. **a**, Workflow of plaNET-seq. Chromatin from a stable NRPB2-FLAG line is isolated and DNase treated. After immunoprecipitation and disruption of protein complexes, RNAPII-attached RNA is purified and used for library construction. The base at the 3’-end of the sequenced RNA is the last base added by the RNAPII complex and therefore aligns to the genomic position of transcriptionally engaged RNAPII. **b**, An example of plaNET-seq coverage profile for the gene At1g25550. Positions of RNAPII are shown for sense (blue) and antisense (red) strands. For comparison, mock-IP (negative control) plaNET-Seq sample, as well as stranded RNA-seq, TSS-seq (Transcription start site sequencing) and DR-seq (Direct RNA sequencing) tracks are also shown. The DR-Seq track reveals sites of mRNA cleavage and polyadenylation (PAS). **c**, Definition of novel transcripts detected by plaNET-seq. Divergent transcripts initiate no more than 500 bp upstream of a coding transcript TSS. Upstream transcripts initiate on the sense strand and partly overlap with an annotated transcript. Convergent transcripts initiate from the 5’-half of a coding gene body on the antisense strand. PAS-associated transcripts initiate from the 3’-half or no more than 20% downstream of its length on the antisense strand. Downstream transcripts initiate within a gene on the sense strand and continue beyond the annotated PAS. Distal antisense transcripts overlap with annotated gene on the antisense strand but initiate further downstream than 20% of the gene’s length. Finally, if a transcript was not described by any of the above mentioned classes, it was defined as an intergenic transcript. **d**, Bar chart of the number of transcripts that fall into the classes described in **(a)**. Known non-coding transcripts in Araport11 are shown in checkered fill and novel transcript identified by plaNET-seq without fill.

### Data analysis

The first 4 bases of both R1 and R2 reads in plaNET-Seq are Unique Molecular Identifiers (UMIs). They were trimmed from read sequences and appended to read names using UMI-Tools v0.5.3. After UMI trimming, the 5’-terminal base of R2 corresponds to the 3’-end of original RNA molecule and thus denotes the genomic position of RNAPII active center. R2 reads were aligned to TAIR10 genome assembly using STAR v2.5.2b in transcriptome-guided mode with the following settings: --outSAMmultNmax 1 --alignEndsType Extend5pOfRead1 --clip3pAdapterSeq GATCGTCGGACT. Ensembl Plants release 28 was used as the source of transcript annotation for alignment. The BAM files were sorted using Samtools v1.3.1. The following categories of reads were filtered out: i) PCR duplicates (UMI-Tools); ii) Reads aligned within 100 bp from any rRNA, tRNA, snRNA or snoRNA gene from Araport11 on either strand (BEDtools v2.17.0); iii) Reads aligned with MAPQ < 10 (Samtools). The filtered BAM files were imported into R environment v3.5.1 using GenomicAlignments_1.18.1 library. The strand orientation of reads was flipped to restore strandness of the original RNA molecules. 3’-terminal bases of flipped reads were found to overlap with 5’ or 3’ splice sites much more frequently than could be expected by chance. Such reads most likely represent splicing intermediates due to co-immunoprecipitation of the spliceosome together with FLAG-tagged RNAPII complexes. These reads were filtered out by overlap with the union of splice sites obtained from both Ensembl Plants 28 (TxDb.Athaliana.BioMart.plantsmart28 package) and Araport11 annotations. In addition, all split reads were removed as possible mature RNA contaminations. The remaining reads are expected to represent the nascent RNA population. Their genomic coverage was exported as strand-specific BigWig and bedGraph files using rtracklayer_1.42.2. The full pipeline is provided in the 01-Alignment_plaNET-Seq.sh and 02-Postprocessing_plaNET-Seq.R scripts in the code repository.

A few existing datasets were remapped in this study. They include pNET-Seq(14) (GSE109974), strand-specific RNA-Seq from(26) (GSE81202), as well as Direct RNA sequencing (DR-Seq) data from(27) (ERP001018) and(28) (ERP003245). The pNET-Seq libraries were processed using our plaNET-Seq pipeline (see above). Remapping of RNA-Seq and DR-Seq data is described in 03-Alignment_GRO-Seq_RNA-Seq_DR-Seq.sh. We also re-used our TSS-Seq data originally published in(29) (GSE113677). Moreover, we used nucleosome occupancy tracks and nucleosome coordinates available from the PlantDHS database(30).

Araport11 annotation was used throughout all further steps of data analysis because it is more comprehensive in terms of non-coding transcripts than both TAIR10 and Ensembl Plants 28 annotations. We adjusted gene borders from Araport11 using TSS-Seq and DR-Seq data. If multiple TSS or PAS tag clusters were connected to the same gene, the strongest of them was chosen as the new border. The relevant code is available in 04-Adjustment_Araport11.R script.

To draw metagene plots of plaNET-Seq and other datasets mentioned above, we merged biological replicates and normalized the tracks to 1 million reads in nuclear protein-coding genes. The X axes of metagene plots represent the genomic intervals of choice which were scaled to the defined number of bins. Intervals overlapping multiple annotated transcription units were excluded from consideration. In particular, both introns and exons were trimmed by 5 bp each side prior to scaling to avoid possible artifacts. The Y axes show the sequencing coverage averaged between the genomic intervals. The code required to reproduce metagene plots from bedGraph tracks is available in 05-Metagenes.R script.

Transcript borders were called *de novo* from each plaNET-Seq sample using groHMM package(31). Intervals which have less than 50% reciprocal overlap on the same strand with any known transcription unit in Araport11 were considered as novel (previously unannotated) transcripts. The novel transcripts were clustered between plaNET-Seq samples and merged to obtain a non-redundant set (n = 7228). They were further classified into divergent, convergent, PAS antisense, distal antisense or intergenic (for more details, see 06-groHMM_pipeline.R).

Differentially transcribed known genes and novel transcripts were called by DESeq2(32) from unnormalized plaNET-Seq tracks with FDR < 0.05 and log2FC > 1 (see 07-DESeq2_pipeline.R)

To calculate the read-through (RT) length, we considered strongly transcribed genes (plaNET-seq FPKM in WT samples above 5). Genomic intervals for RT length estimation were defined from PAS of the analyzed gene to the nearest downstream TSS. Coordinates of TSS and PAS clusters were called from TSS-seq and Direct RNA-seq datasets as described above. For each gene of interest, the empirical distribution of plaNET-seq tag counts in 100 bp sliding window was obtained (the “transcription” model). The “random” model corresponding to the untranscribed state was represented by Poisson distribution where the rate parameter was estimated from plaNET-seq tag counts in intergenic regions. Then PlaNET-seq tags were counted in every 100 bp window moving in 10 bp steps along the candidate RT genomic interval. For each window, the probability to observe at most this tag count under the gene-specific “transcription" model was divided by the probability to observe at least this tag count under the alternative “random” model. The start position of the first window where the probability ratio dropped below 1 was considered as the end of the read-through region. The code is available in 08-Readthrough_distance.R

To calculate promoter-proximal RNAPII stalling index for each gene longer than 1 Kb, we first found 100 bp windows with the highest plaNET-Seq coverage within the interval [TSS − 100 bp, TSS + 300 bp]. Center of this window was considered as the summit of promoter-proximal RNAPII peak. The stalling index was then calculated as the ratio of plaNET-Seq coverage in this window vs the whole gene (normalized by gene width). Similarly, the intronic stalling index (ISI) was calculated for each intron longer than 50 bp: first we found the “best” 10 bp window within the intron, and then we divided its plaNET-Seq coverage by width-normalized coverage of the whole intron. Introns with FPKM-normalized plaNET-Seq coverage above 10 were further classified by their stalling index into “strong” (ISI ≥ 5.5), “medium” (3.5 < ISI < 5.5) and “weak” (ISI ≤ 3.5). For more detailed description, refer to 09-Stalling_index.R.

Chromatin states were downloaded from the PCSD database(33). Based on relative enrichment of different states along protein-coding genes, we combined the original 36 states into 5 groups: “Promoter” (states 13 and 15-21), “Promoter to early elongation” (states 22 and 23), “Early elongation” (states 24-26), “Late elongation” (states 3-12 and 27-28) and “Termination” (states 1 and 2).

## RESULTS

### plaNET-seq robustly detects nascent RNAPII transcription in *Arabidopsis*

To purify RNAPII complexes, we relied on a FLAG-immunoprecipitation of the second-largest RNAPII subunit (NRPB2-FLAG). The NRPB2-FLAG construct covers lethal null-alleles of *nrpb2*, which makes these lines suitable to capture RNAPII as all complexes carry the tagged NRPB2 subunit(24). We used the nuclear fraction of flash-frozen *Arabidopsis* seedlings as starting material (Fig. 1a). RNAPII complexes were immunoprecipitated with high efficiency (Supplementary fig. 1a), and nascent RNA was purified and used for library construction (Supplementary fig. 1b). Processed reads were aligned to the *Arabidopsis* genome, identifying positions of the nascent RNA 3’-ends (Fig. 1b, upper panel). Visualized in a genome browser, plaNET-seq shows the characteristic “spiky” pattern that represents the nascent RNAPII transcription at each nucleotide. Our plaNET-seq libraries showed high reproducibility between replicates and confirmed low-velocity nascent RNAPII transcription at gene boundaries (Supplementary fig. 1c-d). We also generated a mock-IP plaNET-seq library to assess the stringency of our protocol. The signal of mock-IP plaNET-seq libraries was extremely low, supporting FLAG-IP specific signal corresponding to nascent RNAPII transcription in our samples (Fig. 1b, Supplementary fig. 1e). The libraries of nascent RNA appeared enriched for intronic reads and reads downstream of the annotated poly-(A)-site that represented RNAPII complexes undergoing termination of transcription. Steady-state methods such as RNA-seq do not provide this information on nascent RNAPII transcription, further supporting our successful enrichment for nascent RNA (Fig. 1b). We called transcripts *de novo* from plaNET-seq data using the groHMM algorithm(31) and identified thousands of transcripts not annotated in Araport11 (Fig. 1c-d, Supplementary Data 1). The majority of these novel transcripts were in proximity to known genes, or overlapping them on the antisense strand (Fig. 1c-d). Overall, RNA-seq data correlated well with our plaNET-seq data for annotated transcripts but poorly for unannotated transcripts, emphasizing the power of plaNET-seq to capture transcripts undergoing rapid RNA degradation (Supplementary fig. 1f-g).

### Characterization of divergent and convergent transcription

To further characterize the novel transcripts detected by plaNET-seq, we defined transcripts that start upstream (0-500 bp) from the TSS of a protein-coding gene but on the opposite strand as divergent non-coding transcripts (DNC) (Fig. 1c, Fig. 2a). DNC represents an important source of lncRNA transcription in yeast and metazoans(16,34–36), but the presence of DNC in *Arabidopsis* has been questioned(37). plaNET-seq provided evidence for DNC at 917 protein-coding genes and the DNC transcription start site (divTSS) was most often located 200-400 bp upstream from the coding TSS (Fig. 2b). Thus, these data support the presence of DNC in plant genomes, although to a lower extent compared to yeast or mammals. An example of DNC was identified at the At3g28140 locus (Fig. 2c). In general, genes driving DNC in plants had higher nascent RNAPII transcription on the coding strand compared to non-DNC genes (Fig. 2d), indicating that DNC was associated with NDRs of highly expressed genes. Metagene analyses of DNC using TSS-seq data in the *hua enhancer 2-2* mutant (*hen2-2*, a nuclear exosome mutant)(38) showed DNC degradation by the nuclear exosome in *Arabidopsis* (Fig. 2e), similar as in yeast and metazoans(35,39). DNC promoters had higher nucleosome density in the divergent non-coding direction compared to a control set of genes with similar transcription level (Supplementary fig. 2a). DNC promoters exhibited NDRs with well-defined flanking −1 and +1 nucleosomes (Supplementary fig. 2b). In conclusion, DNC transcription shares regulatory principles with budding yeast(40), an association with high definition of the −1 nucleosome, and repressed by co-transcriptional RNA degradation(41).

**Figure 2:**
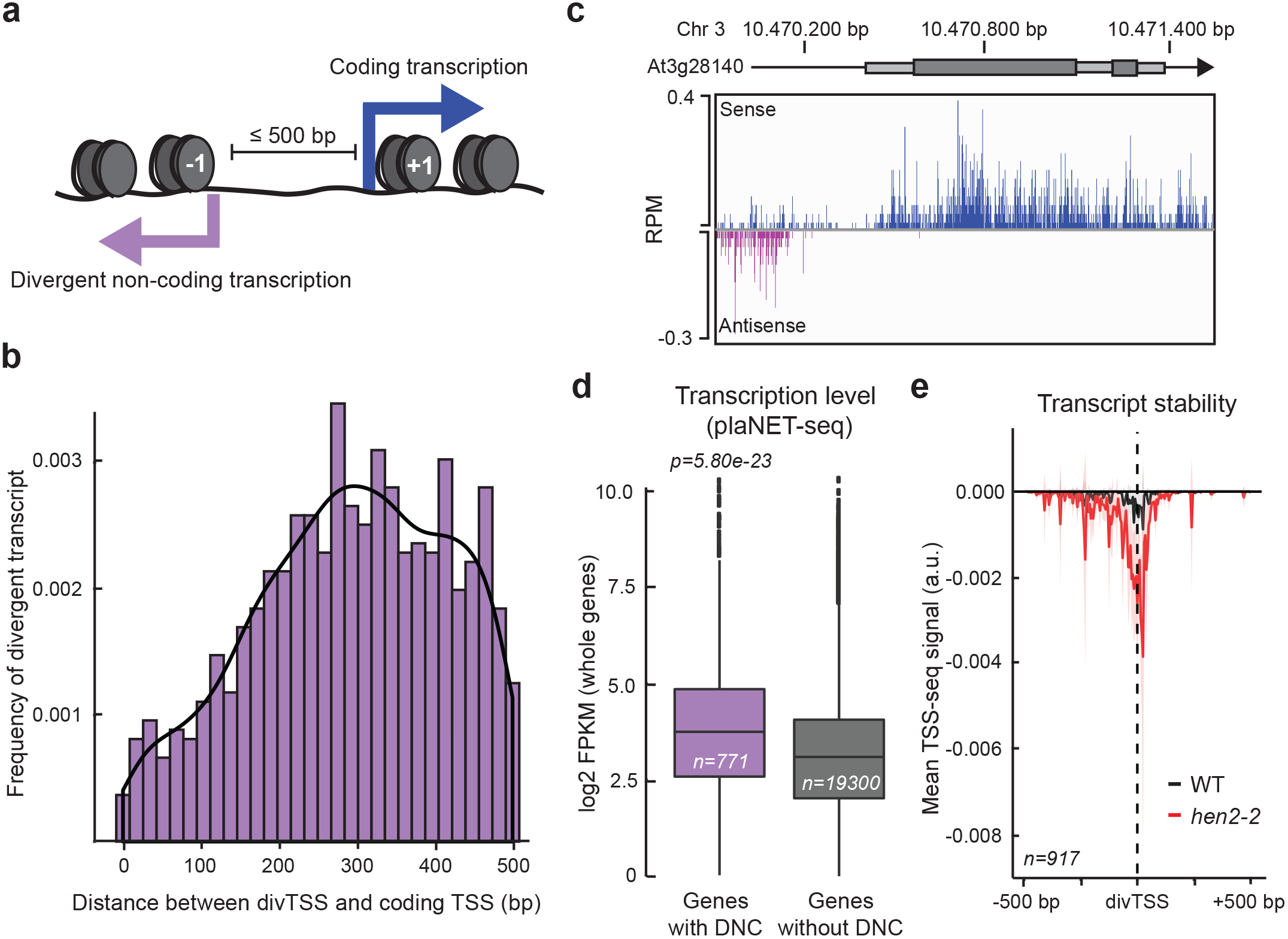
Divergent transcription occurs at highly active NDRs. **a**, Schematic illustration of a divergent promoter. The nucleosomes surrounding the shared NDR are defined as −1 (DNC direction) and +1 (coding direction). **b,** Histogram of the absolute distance between start site for the divergent transcript (divTSS) and the coding TSS (bp). **c,** An example of a divergent promoter (At3g28140). Nascent RNAPII transcription is shown for sense and divergent transcripts in blue and purple, respectively. **d,** Box plot of transcription level of protein-coding genes with a DNC (purple) and without a DNC (grey) as measured by plaNET-seq. Statistical significance of differences was assessed by two-sided Mann-Whitney U test. **e**, Metagene analysis of TSS-Seq signal on the antisense strand of DNC promoters. Wild type signal is shown in black and the nuclear exosome mutant *hen2-2* in red. DNC could be detected with TSS-seq data and DNC were targeted by the nuclear exosome. The shaded area shows 95% confidence interval for the mean.

In addition to DNC, groHMM detected 5313 novel transcripts that overlap a single annotated gene transcription unit fully or partially on the antisense strand (Fig. 3a). We considered novel transcripts as antisense transcripts when they either started internally of a host gene, or no more than 20% of its length downstream (n=4922). We detected two preferential initiation sites for such antisense transcripts along the gene body (Fig. 3a). The predominant peak of initiation site frequency was found at the 3’-end of genes, defined as PAS-associated antisense transcription (n=3223). The second peak was located within the first 50% of the gene body, and we defined these transcripts as convergent antisense transcripts (CAS; n=1699). CASs have been detected in human cells(13,19) but have so far been uncharacterized in plants. The TSS of convergent transcripts (casTSS) most often initiated at a distance between 250 to 1000 bp from the sense TSSs (Supplementary fig. 3a), exemplified by the At2g46710 gene (Fig. 3b). Interestingly, casTSSs showed a strong bias towards the exon-intron boundaries with a peak very close to the 5’ splice sites (5’SS, Fig. 3c). The nucleosome density upstream of the casTSS showed a sharp decrease, suggesting an intragenic NDR (Supplementary fig. 3b). Interestingly, when we assigned previously described chromatin states(33) to the bodies of *Arabidopsis* genes and explored where CAS transcription initiated, we detected an over-representation of casTSS within the chromatin states we denoted as promoter-to-early elongation (Supplementary fig. 3c-d). This indicated that the CAS initiation region coincided with a location where RNAPII complexes enter productive elongation. Genes giving rise to CAS had higher sense strand transcription compared to genes without detectable CAS (Fig. 3d). These data indicated an association of CAS with a subset of highly transcribed genes. In addition, a comparison of TSS-seq data in wild type Col-0 seedlings and *hen2-2* mutants showed that CAS transcripts are nuclear exosome targets (Fig. 3d). Thus, we characterized *Arabidopsis* CAS as nuclear exosome targets that initiate from a NDR in promoter-proximal intervals of highly expressed genes. All in all, our plaNET-seq data highlights the strength of a nascent RNA detection method to identify cryptic non-coding transcripts.

**Figure 3:**
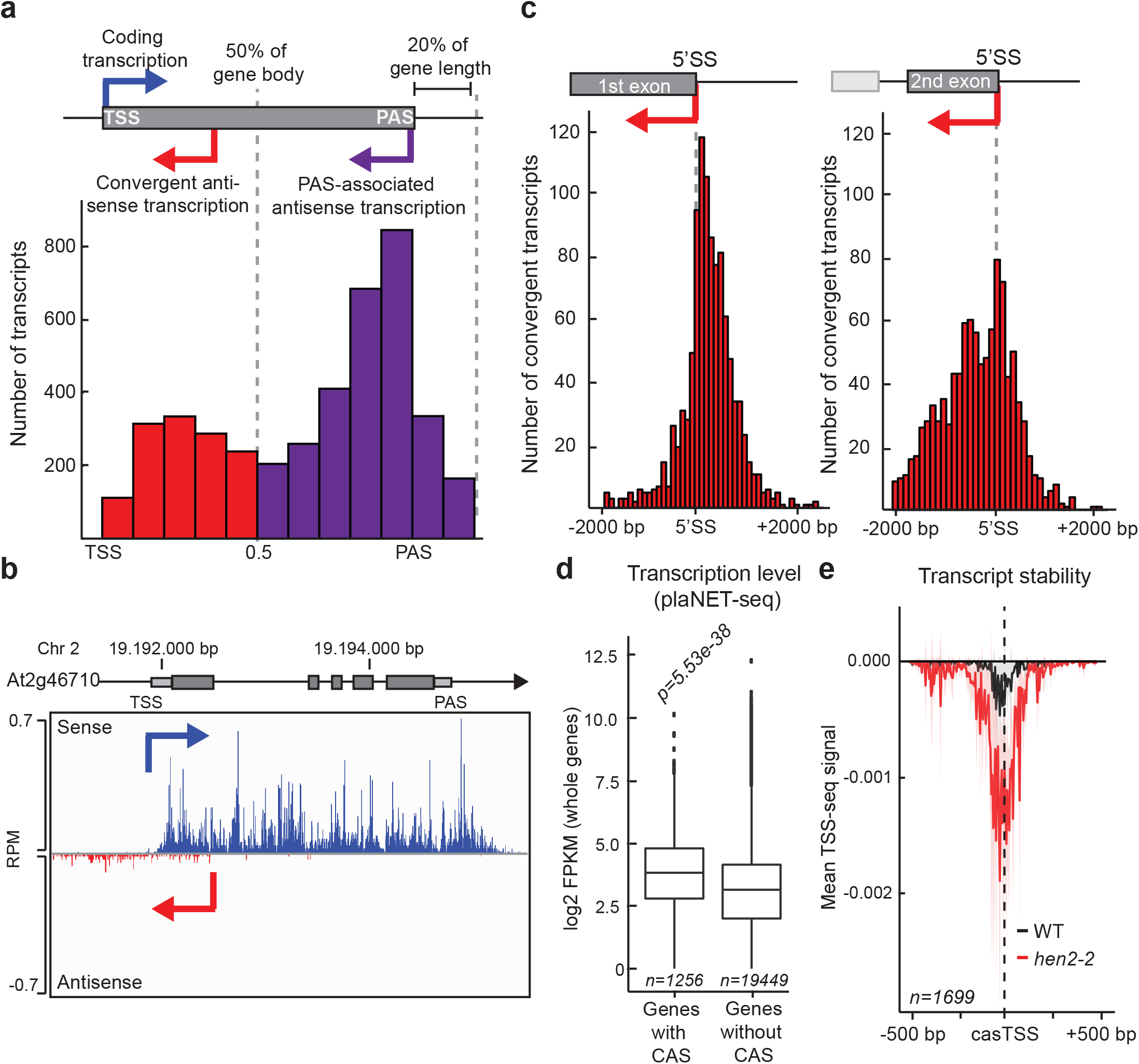
Convergent antisense transcription is a common feature in *Arabidopsis*. **a,** Histogram of the relative distance between initiation sites of antisense transcripts and the sense TSS (expressed as fraction of the sense gene length). Antisense transcription was defined either as convergent (if initiated within the first 50% of the sense gene length: red bars), or as PAS-associated (if initiated within the second 50% of the sense gene length or after the PAS up to a distance of 20% of the gene length after the gene end). **b**, An example of a convergent transcript (At2g46710). Nascent RNAPII transcription is shown for sense and convergent transcripts in blue and red, respectively. **c**, Histogram of distances between the start sites of convergent transcripts and the first 5’ splice site (5’SS) (left panel) or the second 5’SS (right panel). **d,** Box plot of transcription level of coding transcripts with a CAS and without a CAS. Statistical significance of the difference was measured by two-sided Mann-Whitney U test. Genes with a CAS showed higher transcription in the sense direction compared to those without a CAS. **e**, Metagene analysis of TSS-seq signal on the antisense strand in 1 kb windows anchored at the casTSS. Wild type signal is shown in black and the nuclear exosome mutant *hen2-2* in red. CAS could be detected with TSS-seq data, and CAS are targeted by the nuclear exosome. The shaded area shows 95% confidence interval for the mean. **d,** Metagene analysis of nucleosome density in 1 kb windows centered at the convergent transcript start site (casTSS). The shaded area shows 95% confidence interval for the mean. **d**, Metagene analysis of chromatin states determined by ChromHMM along the gene bodies of Arabidopsis genes. Based on the PCSD database, the following states were assigned to respective group: promoter (Prom; states 13, 15-21), promoter-to-early elongation (PromToEarly; states 22-23), early elongation (Early; states 24-26), late elongation (Late; states 3-12, 27-28) and polyA (pA; states 1-2). Each CAS was assigned a chromatin state group based on overlap with casTSS. Observed frequencies of casTSS were plotted together with the expected frequencies of overlap based on the random model. **e**, Metagene analysis of plaNET-Seq signal on the sense and the antisense strand in windows anchored at the casTSS. 22°C (control sample) is shown in black, 3h 4°C in blue, 12h 4°C in light blue. The shaded area shows 95% confidence interval for the mean.

### Low temperature lead to major re-programming of nascent RNAPII transcription

In addition to the capture of cryptic transcripts, NET-seq interrogates the RNAPII transcription dynamics over coding and non-coding transcription units, revealing regions of low-velocity transcription. The link between temperature and transcriptional output in plants(3) lead us to hypothesize that chilling temperatures may regulate nascent RNAPII transcription over these regions. Therefore, we exposed seedlings to early stages of cold-acclimation (3 and 12 hours at 4°C, Fig. 4a). Numerous transcripts had significantly changed plaNET-seq signal over their transcription units in our conditions (Fig. 4b, Supplementary Dataset 2). The number of differentially transcribed known genes at 3h at 4°C versus 22°C greatly exceeded those detected as differentially expressed in the same conditions and identical cut off values by Transcription Start Site sequencing (TSS-seq)(2). These data suggest that the detection of steady-state levels of RNA species (i.e. by TSS-seq) does not fully capture the actual changes in nascent transcription during exposure to 4°C (Fig. 4c)(2). Strikingly, 47% and 50% of known transcripts which were upregulated or downregulated after 3h at 4°C, returned to baseline levels after 12h at 4°C (Fig. 4d), suggesting transient re-programming of nascent RNAPII transcription. Nascent transcription of the novel non-coding transcripts was also affected by the cold treatment, as shown on metagene plots for divergent, convergent and PAS-associated antisense transcripts (Fig. 5a-c). We detected a rapid decrease of plaNET-seq signal after 3h at 4°C that reverted back to or close to control levels after 12h at 4°C. Thus, our results support the notion that transcription of many non-coding transcripts respond rapidly to a changing environment(42). Taken together, plaNET-seq detected genome-wide transcriptional changes with increased sensitivity compared to steady-state methods and revealed a major re-programming of nascent RNAPII transcription in response to chilling temperatures.

**Figure 4:**
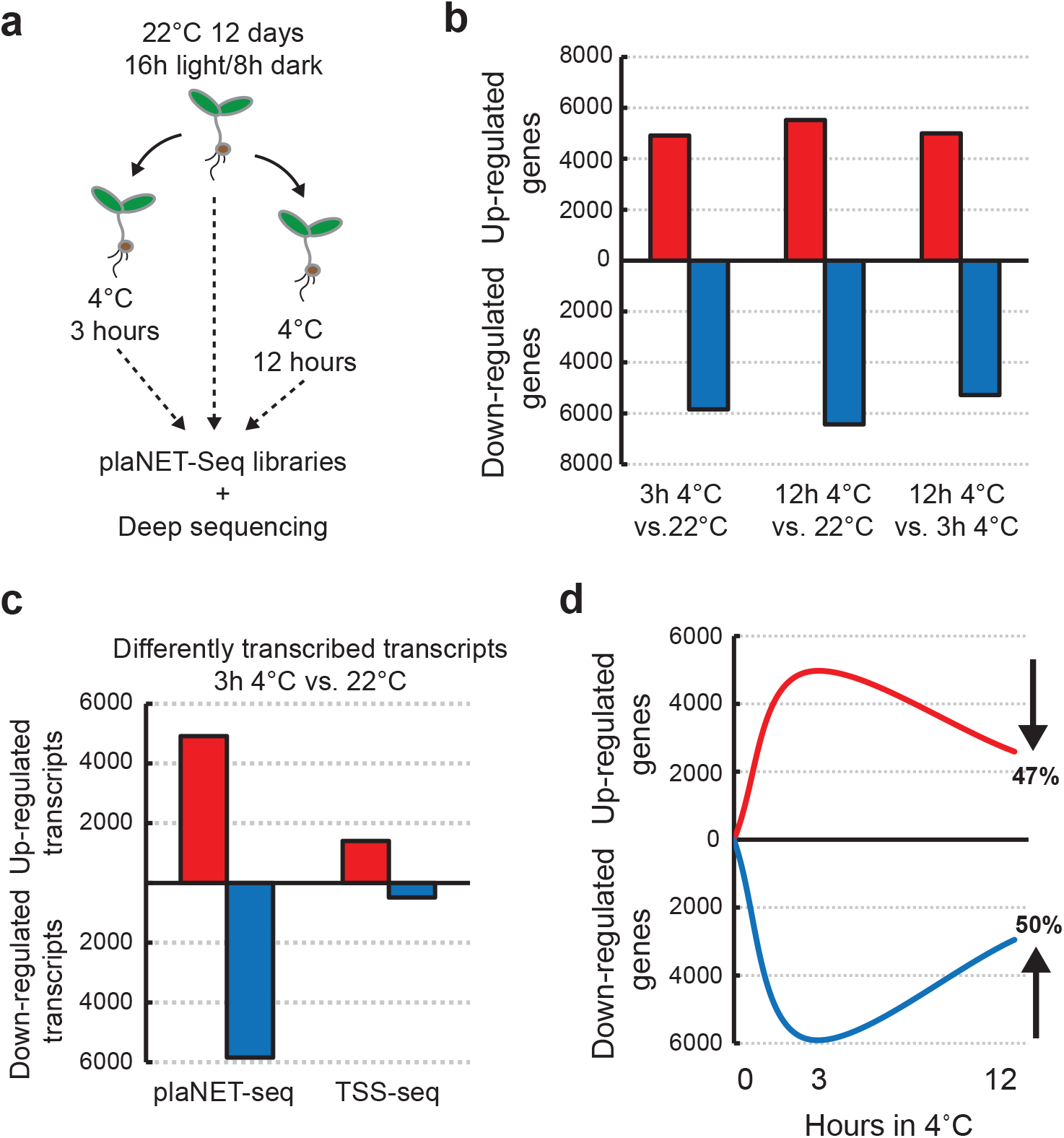
Low temperature leads to re-programming of nascent RNAPII transcription. **a**, Illustration of the experimental design of low temperature exposure. Seedlings were grown for 12 days under a long day light regime on agar plates. Exposure to low temperature was performed for 3 or 12 hours during the light hours and samples collected and flash frozen in liquid nitrogen. **b**, The number of differently transcribed genes determined by plaNET-seq in response to low temperature treatment. **c,** Numbers of up-and down-regulated transcripts after 3 h at 4°C (compared to the control grown at 22°C) as determined by DESeq2 using plaNET-seq and TSS-seq data. The transcriptional changes detected by plaNET-seq exceeded those detected with the same cutoff values by TSS-seq. **d**, Schematic time course of how differently transcribed genes after 3h at 4°C are regulated at 12h at 4°C compared to control.

**Figure 5:**
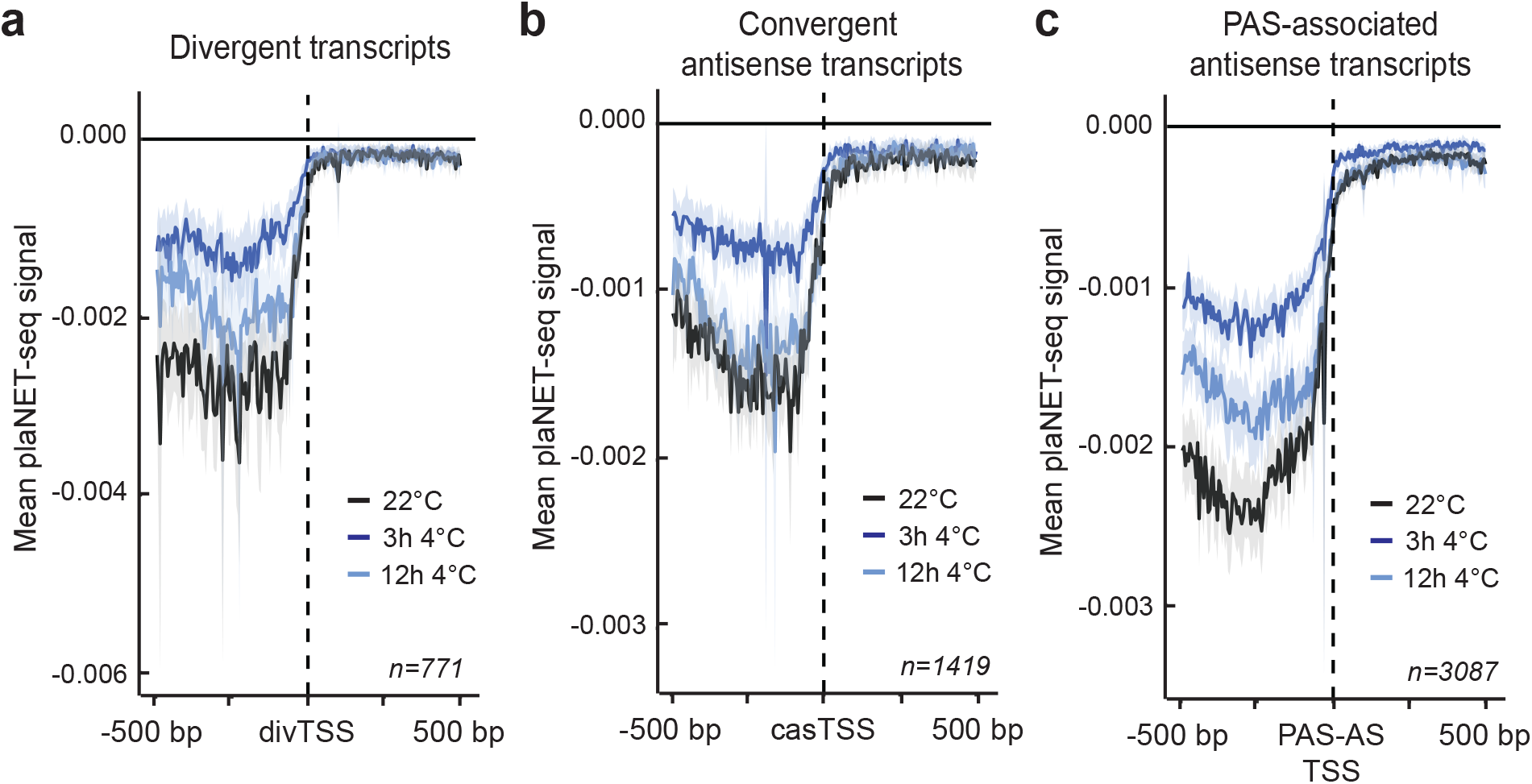
Nascent RNAPII transcription of non-coding transcripts is affected by cold. Metagene analysis of the plaNET-Seq signal in a 1 kb window for **(a)** DNC, anchored at the divTSS, **(b)** CAS, anchored at the casTSS and **(c)** PAS-AS, anchored at the PAS-AS TSS. 22°C (control sample) is shown in black, 3h 4°C in blue, 12h 4°C in light blue. The shaded area shows 95% confidence interval for the mean.

### Exons and co-transcriptional splicing represent transient transcriptional barriers at low temperature

The re-programming of nascent RNAPII transcription in response to chilling temperatures prompted us to look closer at the effects on coding regions in the genome. Eukaryotic genes have exon-intron architecture where introns are co-transcriptionally spliced out to form a functional mRNA. The close proximity of a transcribing RNAPII complex and the spliceosome is detected with NET-seq(15,21). Splicing intermediates can readily be detected in NET-seq data, in particular the 5’ splice site (5’SS) that is protected by the co-purified spliceosome (Fig. 6a), as previously reported in human NET-seq(15). We thus filtered out these read positions in our analysis since the RNAPII-associated RNA 3’-ends through co-purification of the spliceosome may not precisely inform on the position of nascent RNAPII transcription(15). Interestingly, when we plotted the fraction of 5’SS reads in our low temperature exposed plaNET-seq samples, we detected a strong decrease of 5’SS reads after 3h at 4°C compared to 22°C (Fig. 6b). The decrease reverted back to control levels after 12h at 4°C, suggesting that the kinetics of the splicing reaction was initially affected by low temperature (Fig. 6b). Moreover, we detected a transient increase of the exon to intron ratio of nascent RNAPII transcription after 3h at 4°C compared to 22°C and 12 at 4°C (Fig. 6c). These data indicated a transiently increased nascent RNAPII transcription over exons at 4°C. Consistently, many of the transcripts upregulated after 3 hours were relatively long, multi-exonic genes compared to downregulated genes, whereas an inverse relationship was detected for expression changes from 3h to 12h at 4°C (Supplementary fig. 5a-b).

**Figure 6:**
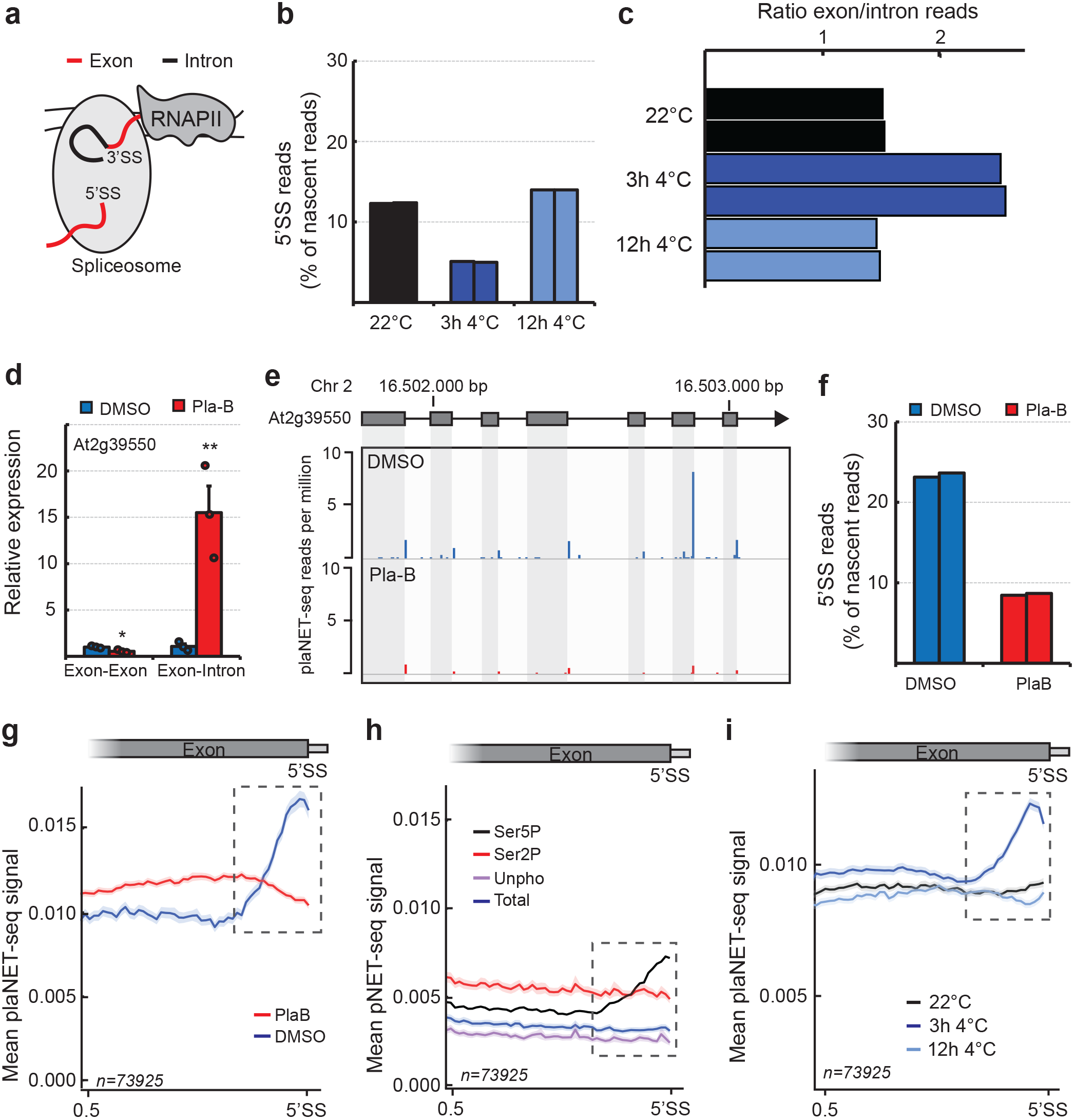
The effect of splicing and intragenic RNAPII stalling. **a**, Illustration of the RNAPII-spliceosome complex during active transcription. The spliceosome protects the 5’SS and the splicing intermediates are co-purified with transcriptionally engaged RNAPII complex in NET-seq. **b**, Bar chart of the percentage of 5’SS intermediates found in the control and low temperature exposed replicates of plaNET-seq. **c**, Histogram showing the ratio between plaNET-seq reads mapping to all exons and all introns in the replicates of low temperature treatment. **d**, RT-qPCR validation of the plaB treatment efficiency (shown for a splicing event of the At2g39550 mRNA). Bars represent mean ± SEM of three biological replicates (circles). The statistical significance of differences was calculated by two-sided t-test. *p<0.05, **p<0.01. **e,** PlaNET-seq co-purifies splicing intermediates, predominantly 5’SS species. The effect of the splicing inhibitor plaB is shown for the gene At2g39550. **f**, Bar chart of the percentage of 5’SS intermediates found in the plaNET-seq DMSO and plaB replicates. **g**, Metagene analysis of nascent RNAPII transcription over the 3’-half of internal exons as determined by plaNET-seq. DMSO is shown in blue and plaB in red. Dashed box indicates stalling site at the 3’-end of exons. The shaded area shows 95% confidence interval for the mean. **h,** Metagene analysis of nascent RNAPII transcription over the 3’-half of internal exons as determined by pNET-seq. Data from the Ser5P antibody is shown in black, Ser2P in red, Unphosphorylated in purple and total RNAPII in blue. Dashed box indicates stalling site at the 3’-end of exons. The shaded area shows 95% confidence interval for the mean. **i**, Metagene analysis of nascent RNAPII transcription over the 3’-half of internal exons as determined by plaNET-seq. 22°C (control sample) is shown in black, 3h 4°C in blue, 12h 4°C in light blue. Dashed box indicates stalling site at the 3’-end of exons. The shaded area shows 95% confidence interval for the mean.

The hypothesis that splicing kinetics may be transiently affected by low temperature prompted us to examine the connection between splicing and RNAPII transcription more closely. We applied the splicing inhibitors pladienolide B (plaB) and herboxidiene and confirmed their effect on sensitive splicing events(43) with RT-qPCR (Fig. 6d, Supplementary fig. 4a). Next, we treated seedlings with DMSO or plaB for 6 hours and generated plaNET-seq libraries. We detected a large decrease in 5’SS reads in our plaB samples compared to the DMSO samples, confirming a successful inhibition of the splicing reaction (Fig. 6e-f). Our analysis identified small nuclear RNAs involved in splicing, confirming co-purification of the spliceosome with RNAPII complexes (Supplementary fig. 4b), consistent with earlier reports(15,21). Metagene profiles of internal exons revealed increased nascent RNAPII transcription upstream of the 5’SS in DMSO compared to plaB, supporting splicing-dependent RNAPII stalling before the end of exons (Fig. 6g, dashed box). This exonic RNAPII stalling was visible also in the re-analyzed pNET-Seq data(14), however only in the serine-5 phosphorylation (Ser5P) track which corresponds to NRPB1 phosphorylated at Ser5 position of its C-terminal domain (Fig. 6h). In our cold-treated samples, we detected an increased peak at the end of exons after 3 hours 4°C compared to 22°C (Fig. 6i, dashed box). The increased height of the peak was transient and reverted to baseline levels after 12h at 4°C. In conclusion, our analyses support a splicing-dependent dynamic increase of nascent RNAPII transcription at the end of exons during low temperature. These data may indicate that the kinetics of the splicing reaction is transiently reduced in the chilling response.

### Identification of a novel intragenic RNAPII stalling site

In introns, plaNET-Seq metagene profiles of our plaB and DMSO samples revealed a peak of nascent RNAPII transcription close to the 5’SS (Supplementary fig. 6a). Moreover, this intronic peak is most clearly visible in the Ser5P track of pNET-Seq data (Supplementary fig. 6b). We called the peak coordinates in each intron using sliding window approach on Ser5p pNET-Seq data. Next, we calculated an “Intronic stalling index” (ISI) for each intron based on the plaNET-Seq data in untreated Col-0 sample. Finally, we divided the introns based on ISI into those with strong, medium or weak stalling (for more details, see Methods). The intronic peak was most frequently observed at 25 nt downstream of the 5’SS, irrespective of the ISI level (Fig. 7a). Grouping introns by ISI revealed that introns with higher ISI scores were on average longer than low ISI-score introns (Supplementary fig. 6c). We therefore stratified introns according to their length to explore potential effects of the intronic peak. We detected no evidence for increased nucleosome signal in short introns (60-250 bp (*n=97558*), Supplementary fig. 6d). However, we detected peaks in nucleosome density in longer introns (250-1000 bp, *n=15991*), suggesting that these included one or several phased nucleosomes (Supplementary fig. 6d). We next plotted nascent RNAPII transcription over long introns compared to a control set of short introns (obtained from the same genes to avoid any effect of gene expression level). We detected a higher plaNET-Seq signal over longer introns, suggesting that long introns were transcribed more slowly compared to short introns (Supplementary fig. 6e). Thus, nucleosome barriers may contribute to a reduced transcription speed and increased plaNET-seq signal of longer introns. Interestingly, the intronic peak in short introns was largely plaB insensitive (Fig. 7b), whereas stalling in long introns was sensitive to plaB (Fig. 7c). Similarly, our cold-treated samples showed small effects of the intron peak for short introns (Fig. 7d) but a large increase of nascent RNAPII transcription after 3h at 4°C that reverted back to control levels after 12h at 4°C in long introns (Fig. 7e). This observation further supported a transient decrease in kinetics of the splicing reaction after low temperature exposure. All in all, our plaB and cold-treated samples provide key information to distinguish plaNET-seq signal that is dependent on the splicing reaction from peaks of RNAPII activity independent of splicing. Our data support a RNAPII stalling site 25 nt into plant introns. The sensitivity of this peak to plaB and to low temperature correlates with intron length, perhaps indicating RNAPII-stalling associated checkpoint to improve splicing accuracy of long introns. The intronic peak of RNAPII activity represents a novel site of RNAPII stalling during gene transcription that represents the 3^rd^ stalling site in addition to the positions at gene boundaries.

**Figure 7:**
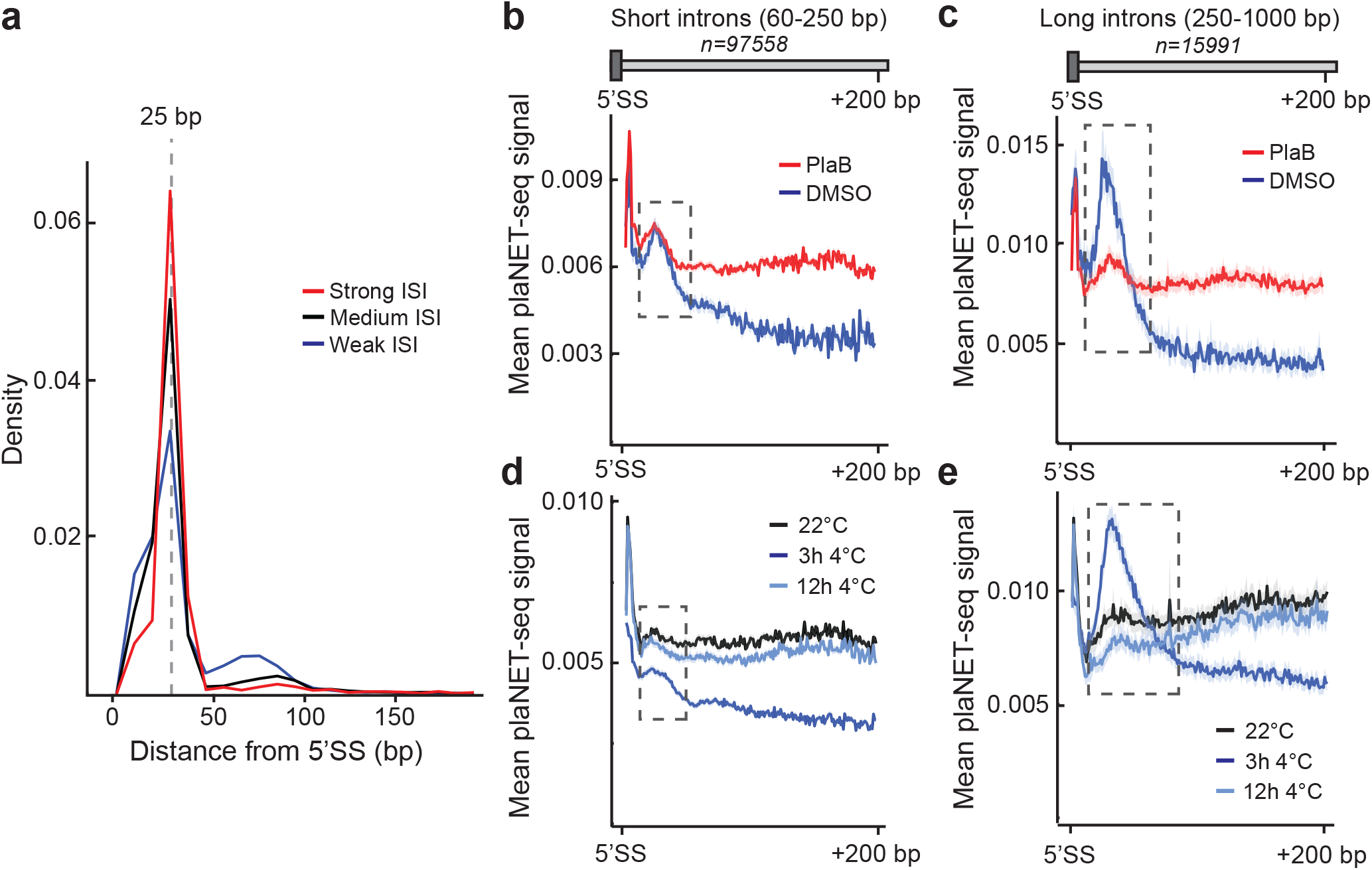
Identification of a novel RNAPII stalling site in introns. **a,** Absolute distance of the intronic peak from the 5’SS. Only introns with FPKM-normalized plaNET-Seq coverage above 10 are shown. Introns with strong intronic stalling index (ISI ≥ 5.5) are shown in red, medium (3.5 < ISI < 5.5) in black and weak (ISI ≤ 3.5) in blue. **b**, Metagene analysis of nascent RNAPII transcription in short introns as determined by plaNET-Seq. DMSO is shown in blue and plaB in red. Dashed box indicates stalling site at the 3’-end of exons. The shaded area shows 95% confidence interval for the mean. **c,** Metagene analysis of nascent RNAPII transcription in long introns as determined by plaNET-Seq. DMSO is shown in blue and plaB in red. Dashed box indicates stalling site at the 3’-end of exons. The shaded area shows 95% confidence interval for the mean. **d,** Metagene analysis of nascent RNAPII transcription in short introns as determined by plaNET-Seq. 22°C (control sample) is shown in black, 3h 4°C in blue, 12h 4°C in light blue. Dashed box indicates stalling site at the 3’-end of exons. The shaded area shows 95% confidence interval for the mean. **e,** Metagene analysis of nascent RNAPII transcription in long introns as determined by plaNET-Seq. 22°C (control sample) is shown in black, 3h 4°C in blue, 12h 4°C in light blue. Dashed box indicates stalling site at the 3’-end of exons. The shaded area shows 95% confidence interval for the mean.

### Low temperature effects promoter-proximal RNAPII stalling

To further investigate RNAPII stalling at gene boundaries, we first focused on the beginning of transcription units (i.e. promoter-proximal stalling). plaNET-seq detected a large fraction of reads at 5’-ends of genes, consistent with previous studies in plants and metazoans(5,14) (Supplementary fig. 1c). We found no clear correlation between the annotated TSS position and the maximal density of nascent RNA signal on the sense strand (Supplementary fig. 6f). To test if other genomic features could offer a better correlation we used nucleosome positioning data (MNase-seq). Metagene plots anchored at the center of the first nucleosome revealed a strong association with peaks of nascent RNAPII transcription (Fig. 8a), suggesting a nucleosome defined promoter proximal stalling mechanism in *Arabidopsis*. Metagene profiles for 0, 3 and 12 hours at 4°C indicated that low temperature affected RNAPII stalling at the first (i.e. +1) nucleosome (Fig. 8a). 3h at 4°C resulted in an increased peak around the center of the +1 nucleosome, indicating greater promoter-proximal stalling. In contrast, the 12h 4°C samples resulted in decreased stalling. These results prompted us to investigate if pools of RNAPII engaged in promoter-proximal stalling may facilitate temperature-dependent gene regulation. We calculated a “Promoter-proximal stalling index” from plaNET-seq data (i.e. relative nascent RNAPII transcription at the promoter proximal region versus the gene body) as previously described(14). Transcripts that were up-regulated after 3h at 4°C showed a significantly increased stalling index before low temperature treatment (22°C). In addition, transcripts that were down-regulated after 3h at 4°C exhibited significantly decreased promoter proximal stalling compared to non-regulated transcripts (Fig. 8b). These results support a role for RNAPII promoter-proximal stalling to adjust transcription to low temperature. In conclusion, plaNET-seq revealed a nucleosome defined promoter-proximal RNAPII stalling mechanism that may facilitate reprogramming of gene expression in response to temperature changes.

**Figure 8:**
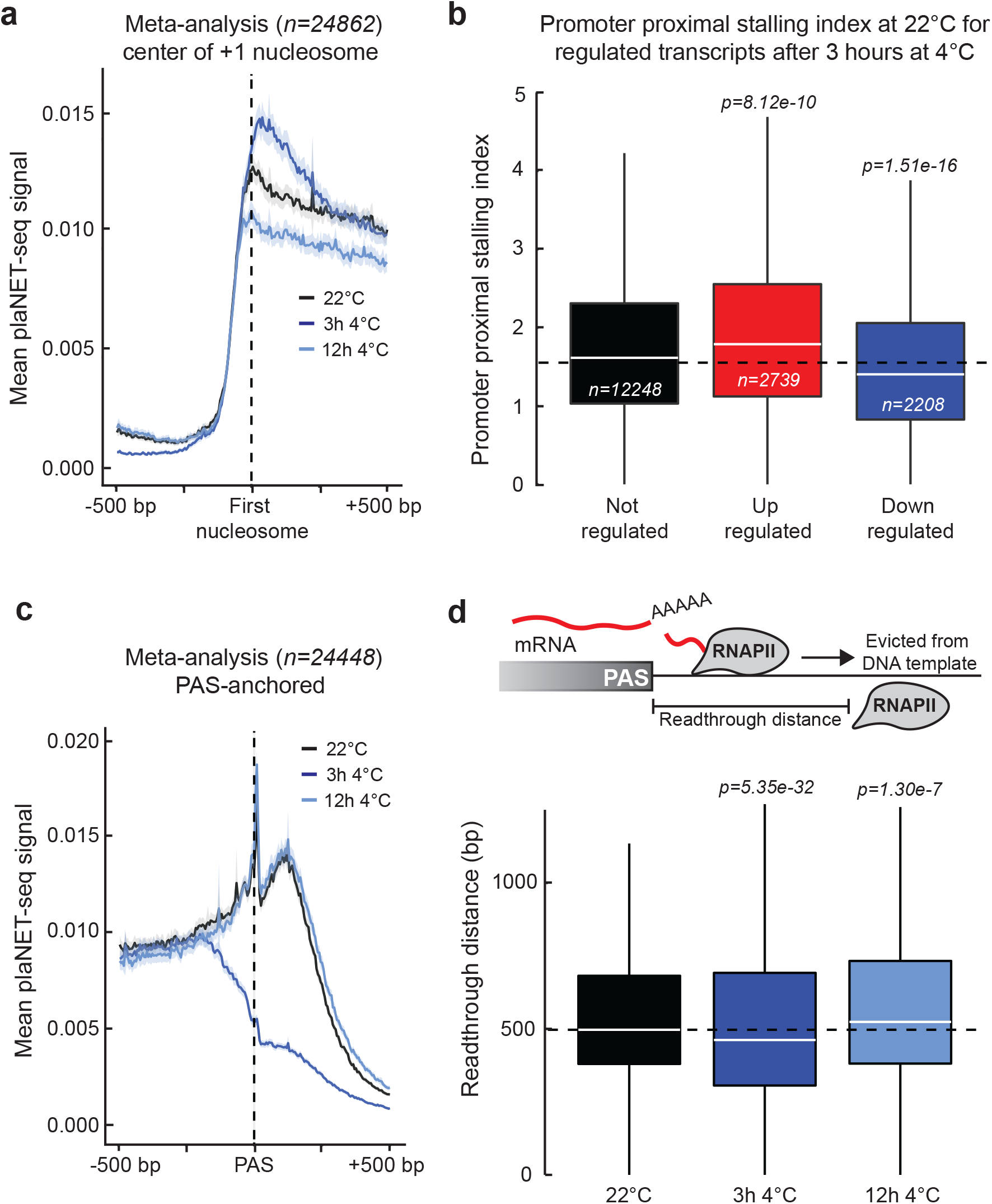
Low temperature affects RNAPII stalling at gene boundaries. **a**, Metagene analysis of the plaNET-Seq signal in a 1 kb window anchored at the center of +1 nucleosome. 22°C (control sample) is shown in black, 3h 4°C in blue, 12h 4°C in light blue. The shaded area shows 95% confidence interval for the mean. **b**, Box plot of promoter-proximal stalling index in control conditions (22°C) of genes which are differently transcribed at 3h 4°C. Black denotes transcripts with unchanged expression, red denotes upregulated transcripts and blue denotes downregulated transcripts. Statistical differences were assessed by two-sided Mann-Whitney U test. **c**, Metagene analysis of the plaNET-Seq signal in a 1 kb window anchored at the PAS. 22°C is shown in black, 3h 4°C is shown in blue, 12h 4°C is shown in light blue. The shaded area shows 95% confidence interval for the mean. **d**, Upper panel illustrates the definition of read-through distance while lower panel shows a box plot of the read-through distance (bp) in 22°C (black), 3h 4°C (blue) and 12h 4°C (light blue) samples. Statistical differences were assessed by two-sided Mann-Whitney U test.

### Low temperature transiently reduces 3’-end associated RNAPII stalling and read-through transcription

In addition to promoter-proximal positions, RNAPII stalls near 3’-ends of *Arabidopsis* genes(14,37,44). We detected increased nascent RNAPII transcription downstream of the poly(A) sites (PAS) (Fig. 8c). We plotted the mean plaNET-seq signal anchored on PAS sites to examine the effect of low temperature on PAS-associated RNAPII stalling. As expected, samples taken before the treatment (22°C) and after 12h at 4°C showed that RNAPII stalled downstream of the PAS (Fig. 8c). Surprisingly, the peak of RNAPII stalled downstream of the PAS was abolished after 3h at 4°C, suggesting a major change in transcription dynamics associated with termination (Fig. 8c). RNAPII complexes transcribe beyond the PAS, representing the zone of transcription termination (Fig. 8d, upper panel). At control conditions (22°C), we detected a median read-through distance of 497 bp (Fig. 8d). This distance was significantly decreased at 3h 4°C (median 462 bp, Fig. 8d). However, at 12h 4°C, we detected a slightly increased read-through distance (median 524 bp, Fig. 8d). Thus, genome-wide distribution of RNAPII such as PAS-associated stalling and read-through distance were transiently altered by low temperature.

## DISCUSSION

### plaNET-seq reveals novel transcription units near annotated genes

Here, we have used NET-seq to study how nascent RNAPII transcription adjusts to low temperature in *Arabidopsis*. Our data detected numerous novel transcripts adjacent and antisense to coding sequences. We identified divergent transcription (DNC) from promoter NDRs in *Arabidopsis*, although at a limited number of genes (Fig. 2) compared to other eukaryotes(15,35). *Arabidopsis* promoters displaying DNC have high expression in the sense direction (i.e. mRNA) and a well-defined NDR with well-positioned −1 and +1 nucleosomes (Fig. 2). However, highly expressed genes in *Arabidopsis* exist without evidence for DNC originating from their promoter NDR. We confirm the repressive effect of nuclear RNA degradation on the detection of DNC. Future studies will be required to elucidate the function of DNC, and the molecular mechanisms that direct RNAPII more strictly into the direction of mRNA transcription at shared promoter NDRs in *Arabidopsis* compared to metazoans. *Arabidopsis* also show extensive antisense initiation from promoter proximal exon-introns boundaries (i.e. CAS; Fig. 3), a common form of antisense transcription in human cells(13,19). In human and plants, promoter proximal introns regulate gene expression and include many *cis*-elements for transcription factor binding(45,46), which may explain the favored site of initiation for CAS. A focused functional dissection of CAS is currently lacking, however CAS transcription may shape the chromatin environment of the corresponding sense promoter as suggested in yeast and human(13,19,47). The casTSSs overlapped frequently with chromatin states which correspond to the transition zone for RNAPII between initiation and productive elongation (Fig. 3), thus highlighting the effects of intragenic chromatin dynamics on TSS selection(29). In summary, our identification of thousands novel transcription units enabled us to detect non-coding transcription linked to gene expression at equivalent positions of transcription units across eukaryotes.

### Co-transcriptional splicing may decrease in response to low temperature

Our results reveal an intragenic peak of RNAPII activity located towards the end of exons (Fig. 6g-i). Exons have well-positioned nucleosomes in human(48) and *Arabidopsis*(49) that may alter RNAPII progression to result in gradual accumulation of nascent RNAPII transcription towards the end of exons. Our data show that the exonic peak is most pronounced after 3h 4°C, perhaps reflecting challenges to transcribe through nucleosome-rich regions during initial low temperature exposure. We detected a similar position of the major stalling site within exons close to the 5’SS in DMSO-treated samples, however this exonic RNAPII stalling was abolished when splicing was chemically inhibited by plaB (Fig. 6g). These data argue that the transient peak of RNAPII at the end of exons may reflect the impact of altered splicing kinetics on nascent RNAPII transcription(50). The decreased RNAPII speed nearby 5’SS may be used by the plant for regulation of alternative splicing events, a biologically essential mechanism for cold acclimation in *Arabidopsis*(23). In addition to the exonic peak, we detect a sharp peak of nascent RNAPII transcription at about 25 bp into introns. This intronic peak co-localized with RNAPII decorated by CTD-Ser5P, a post-translational RNAPII modification that has previously been linked to splicing(15). Interestingly, this peak has not been detected in yeast or human cells, arguing for diverse transcription dynamics within gene bodies between eukaryotes. We identified a reduction of nascent RNAPII transcription at the intronic peak in response to splicing inhibition by plaB treatment for long introns (i.e. 250-1000 bp). These data reveal unprecedented insight into the connections between RNAPII stalling, splicing and intron length that shape plant gene expression. We consider it plausible that this intronic RNAPII peak may represent a checkpoint for accurate splicing of long introns, where we imagine the canonical splice sites to be in a greater competition with cryptic intronic splice sites.

### Low temperature effects RNAPII stalling at gene boundaries

Our analyses of nascent RNAPII transcription highlights the relevance for mechanisms regulation “post-initiation”, in other words beyond RNAPII recruitment to gene promoters through sequence-specific transcription factors. At the 5’-end of genes, RNAPII stalls at the +1 nucleosome during *Arabidopsis* gene expression (Fig. 8a). In human, RNAPII complexes stall at a narrow window of 20-60 bp between the TSS and the +1 nucleosome boundary(15). The stalling in metazoans is influenced by the Negative Elongation Factor (NELF) complex that prevents RNAPII complexes to proceed into productive elongation(51). Interestingly, NELF is conspicuously absent in plants, which may reconcile our identification of the +1 nucleosome as the main determinant for promoter-proximal RNAPII stalling. Our data support the idea that RNAPII complexes stalled at promoter-proximal positions may be released to adjust transcription in response to decreased temperature in *Arabidopsis* (Fig. 8b). A key modulator of temperature-dependent plant gene expression is the histone variant H2A.Z incorporated into the +1 nucleosome(52). It is tempting to speculate that temperature-regulated properties of the +1 nucleosome contribute to temperature-induced expression changes of plant genes by effects on promoter-proximal RNAPII stalling.

At the 3’-end of genes we find a transient chilling-induced contraction of transcription units (Fig. 8c-d). We calculate the read-through distance in *Arabidopsis* to a median length of 497 bp (Fig. 8d). This can be compared to the median read-through distance in *S. cerevisiae* (200 bp)(53) and human cells (3300 bp)(20). The difference in read-through distance may be connected to the level of genome compaction; both *Arabidopsis* and *S. cerevisiae* have gene-denser genomes compared to humans. Gene-dense genomes increase the probability of RNAPII collisions by read-through transcription with harmful consequences for genome stability(54,55). We have not failed to notice that the transient effects on RNAPII read-through distance during low temperature exposure could be consistent with changes in liquid phase viscosity implicated in *Arabidopsis* 3’-end formation(56). Perhaps, our 3h 4°C time-point captures cells during a metabolic adjustment of nuclear liquid environments including those promoting 3’-end formation.

In conclusion, the temperature-induced genome-wide adaptions required to maintain cellular functions provide insight into molecular alterations that promote organismal fitness during environmental change. Our work identifies key parameters of nascent RNAPII transcription that control the transcriptional cold-response in *Arabidopsis* and possibly other eukaryotes.

## AVAILABILITY

The scripts required to reproduce all results and figures are available at GitHub: [https://github.com/Maxim-Ivanov/Kindgren_et_al_2019].

## ACCESSION NUMBERS

plaNET-Seq data is available at NCBI GEO database with accession code GSE131733 (reviewer token: mneleckgrzynjsz).

## SUPPLEMENTARY DATA

Supplementary data consisting of six figures and three data files are available as a separate document.

## ACKNOWLEDGEMENTS

We would like to thank Albin Sandelin and members of the Marquardt lab for critical reading of the manuscript. Adam R. Morris from Bioo Scientific is acknowledged for assistance with modifications to the library construction protocol.

## REFERENCES

1 Markovskaya, E.F. and Shibaeva, T.G. (2017) Low temperature sensors in plants: Hypotheses and assumptions. Biology Bulletin, 44, 150–158.

2 Kindgren, P., Ard, R., Ivanov, M. and Marquardt, S. (2018) Transcriptional read-through of the long non-coding RNA SVALKA governs plant cold acclimation. Nature Communications, 9, 4561.

3 Thomashow, M.F. (1999) PLANT COLD ACCLIMATION: Freezing Tolerance Genes and Regulatory Mechanisms. Annual Review of Plant Physiology and Plant Molecular Biology, 50, 571–599.

4 Mayer, A., Landry, H.M. and Churchman, L.S. (2017) Pause & go: from the discovery of RNA polymerase pausing to its functional implications. Current opinion in cell biology, 46, 72–80.

5 Adelman, K. and Lis, J.T. (2012) Promoter-proximal pausing of RNA polymerase II: emerging roles in metazoans. Nature reviews. Genetics, 13, 720–731.

6 Rougvie, A.E. and Lis, J.T. (1988) The RNA polymerase II molecule at the 5′ end of the uninduced hsp70 gene of D. melanogaster is transcriptionally engaged. Cell, 54, 795–804.

7 Bunch, H., Zheng, X., Burkholder, A., Dillon, S.T., Motola, S., Birrane, G., Ebmeier, C.C., Levine, S., Fargo, D., Hu, G. et al. (2014) TRIM28 regulates RNA polymerase II promoter-proximal pausing and pause release. Nature structural & molecular biology, 21, 876–883.

8 Proudfoot, N.J. (2016) Transcriptional termination in mammals: Stopping the RNA polymerase II juggernaut. Science, 352, aad9926.

9 Connelly, S. and Manley, J.L. (1988) A functional mRNA polyadenylation signal is required for transcription termination by RNA polymerase II. Genes & Development, 2, 440–452.

10 Buratowski, S. (2005) Connections between mRNA 3′ end processing and transcription termination. Current Opinion in Cell Biology, 17, 257–261.

11 Hazelbaker, Dane Z., Marquardt, S., Wlotzka, W. and Buratowski, S. (2013) Kinetic Competition between RNA Polymerase II and Sen1-Dependent Transcription Termination. Molecular Cell, 49, 55–66.

12 Vilborg, A., Sabath, N., Wiesel, Y., Nathans, J., Levy-Adam, F., Yario, T.A., Steitz, J.A. and Shalgi, R. (2017) Comparative analysis reveals genomic features of stress-induced transcriptional readthrough. Proceedings of the National Academy of Sciences, 114, E8362–E8371.

13 Mayer, A., di Iulio, J., Maleri, S., Eser, U., Vierstra, J., Reynolds, A., Sandstrom, R., Stamatoyannopoulos, John A. and Churchman, L.S. (2015) Native Elongating Transcript Sequencing Reveals Human Transcriptional Activity at Nucleotide Resolution. Cell, 161, 541–554.

14 Zhu, J., Liu, M., Liu, X. and Dong, Z. (2018) RNA polymerase II activity revealed by GRO-seq and pNET-seq in Arabidopsis. Nature Plants, 4, 1112–1123.

15 Nojima, T., Gomes, T., Grosso, Ana Rita F., Kimura, H., Dye, Michael J., Dhir, S., Carmo-Fonseca, M. and Proudfoot, Nicholas J. (2015) Mammalian NET-Seq Reveals Genome-wide Nascent Transcription Coupled to RNA Processing. Cell, 161, 526–540.

16 Churchman, L.S. and Weissman, J.S. (2011) Nascent transcript sequencing visualizes transcription at nucleotide resolution. Nature, 469, 368.

17 Schlackow, M., Nojima, T., Gomes, T., Dhir, A., Carmo-Fonseca, M. and Proudfoot, N.J. (2017) Distinctive Patterns of Transcription and RNA Processing for Human lincRNAs. Molecular Cell, 65, 25–38.

18 Andersson, R., Refsing Andersen, P., Valen, E., Core, L.J., Bornholdt, J., Boyd, M., Heick Jensen, T. and Sandelin, A. (2014) Nuclear stability and transcriptional directionality separate functionally distinct RNA species. Nature Communications, 5, 5336.

19 Brown, T., Howe, F.S., Murray, S.C., Wouters, M., Lorenz, P., Seward, E., Rata, S., Angel, A. and Mellor, J. (2018) Antisense transcription-dependent chromatin signature modulates sense transcript dynamics. Molecular Systems Biology, 14, e8007.

20 Schwalb, B., Michel, M., Zacher, B., Frühauf, K., Demel, C., Tresch, A., Gagneur, J. and Cramer, P. (2016) TT-seq maps the human transient transcriptome. Science, 352, 1225–1228.

21 Nojima, T., Rebelo, K., Gomes, T., Grosso, A.R., Proudfoot, N.J. and Carmo-Fonseca, M. (2018) RNA Polymerase II Phosphorylated on CTD Serine 5 Interacts with the Spliceosome during Co-transcriptional Splicing. Molecular Cell, 72, 369–379.e364.

22 Capovilla, G., Delhomme, N., Collani, S., Shutava, I., Bezrukov, I., Symeonidi, E., de Francisco Amorim, M., Laubinger, S. and Schmid, M. (2018) PORCUPINE regulates development in response to temperature through alternative splicing. Nature Plants, 4, 534–539.

23 Calixto, C.P.G., Guo, W., James, A.B., Tzioutziou, N.A., Entizne, J.C., Panter, P.E., Knight, H., Nimmo, H.G., Zhang, R. and Brown, J.W.S. (2018) Rapid and Dynamic Alternative Splicing Impacts the Arabidopsis Cold Response Transcriptome. The Plant Cell, 30, 1424–1444.

24 Onodera, Y., Nakagawa, K., Haag, J.R., Pikaard, D., Mikami, T., Ream, T., Ito, Y. and Pikaard, C.S. (2008) Sex-biased lethality or transmission of defective transcription machinery in Arabidopsis. Genetics, 180, 207–218.

25 Exner, V., Taranto, P., Schönrock, N., Gruissem, W. and Hennig, L. (2006) Chromatin assembly factor CAF-1 is required for cellular differentiation during plant development. Development, 133, 4163–4172.

26 Kohnen, M.V., Schmid-Siegert, E., Trevisan, M., Petrolati, L.A., Sénéchal, F., Müller-Moulé, P., Maloof, J., Xenarios, I. and Fankhauser, C. (2016) Neighbor Detection Induces Organ-Specific Transcriptomes, Revealing Patterns Underlying Hypocotyl-Specific Growth. The Plant Cell, 28, 2889–2904.

27 Sherstnev, A., Duc, C., Cole, C., Zacharaki, V., Hornyik, C., Ozsolak, F., Milos, P.M., Barton, G.J. and Simpson, G.G. (2012) Direct sequencing of Arabidopsis thaliana RNA reveals patterns of cleavage and polyadenylation. Nature Structural & Molecular Biology, 19, 845.

28 Schurch, N.J., Cole, C., Sherstnev, A., Song, J., Duc, C., Storey, K.G., McLean, W.H.I., Brown, S.J., Simpson, G.G. and Barton, G.J. (2014) Improved annotation of 3’ untranslated regions and complex loci by combination of strand-specific direct RNA sequencing, RNA-Seq and ESTs. PLoS One, 10.1371/journal.pone.0094270.

29 Nielsen, M., Ard, R., Leng, X., Ivanov, M., Kindgren, P., Pelechano, V. and Marquardt, S. (2019) Transcription-driven chromatin repression of Intragenic transcription start sites. PLOS Genetics, 15, e1007969.

30 Zhang, T., Zhang, W. and Jiang, J. (2015) Genome-Wide Nucleosome Occupancy and Positioning and Their Impact on Gene Expression and Evolution in Plants. Plant Physiology, 168, 1406–1416.

31 Chae, M., Danko, C.G. and Kraus, W.L. (2015) groHMM: a computational tool for identifying unannotated and cell type-specific transcription units from global run-on sequencing data. BMC Bioinformatics, 16, 222.

32 Love, M.I., Huber, W. and Anders, S. (2014) Moderated estimation of fold change and dispersion for RNA-seq data with DESeq2. Genome Biology, 15, 550.

33 Liu, Y., Tian, T., Zhang, K., You, Q., Yan, H., Zhao, N., Yi, X., Xu, W. and Su, Z. (2017) PCSD: a plant chromatin state database. Nucleic Acids Research, 46, D1157–D1167.

34 Core, L.J., Waterfall, J.J. and Lis, J.T. (2008) Nascent RNA Sequencing Reveals Widespread Pausing and Divergent Initiation at Human Promoters. Science, 322, 1845–1848.

35 Neil, H., Malabat, C., d’Aubenton-Carafa, Y., Xu, Z., Steinmetz, L.M. and Jacquier, A. (2009) Widespread bidirectional promoters are the major source of cryptic transcripts in yeast. Nature, 457, 1038.

36 Seila, A.C., Calabrese, J.M., Levine, S.S., Yeo, G.W., Rahl, P.B., Flynn, R.A., Young, R.A. and Sharp, P.A. (2008) Divergent Transcription from Active Promoters. Science, 322, 1849–1851.

37 Hetzel, J., Duttke, S.H., Benner, C. and Chory, J. (2016) Nascent RNA sequencing reveals distinct features in plant transcription. Proceedings of the National Academy of Sciences, 113, 12316–12321.

38 Lange, H., Zuber, H., Sement, F.M., Chicher, J., Kuhn, L., Hammann, P., Brunaud, V., Bérard, C., Bouteiller, N., Balzergue, S. et al. (2014) The RNA Helicases AtMTR4 and HEN2 Target Specific Subsets of Nuclear Transcripts for Degradation by the Nuclear Exosome in Arabidopsis thaliana. PLOS Genetics, 10, e1004564.

39 Jensen, Torben H., Jacquier, A. and Libri, D. (2013) Dealing with Pervasive Transcription. Molecular Cell, 52, 473–484.

40 Marquardt, S., Escalante-Chong, R., Pho, N., Wang, J., Churchman, L.S., Springer, M. and Buratowski, S. (2014) A Chromatin-Based Mechanism for Limiting Divergent Noncoding Transcription. Cell, 157, 1712–1723.

41 Xu, Z., Wei, W., Gagneur, J., Perocchi, F., Clauder-Münster, S., Camblong, J., Guffanti, E., Stutz, F., Huber, W. and Steinmetz, L.M. (2009) Bidirectional promoters generate pervasive transcription in yeast. Nature, 457, 1033–1037.

42 Matsui, A., Ishida, J., Morosawa, T., Mochizuki, Y., Kaminuma, E., Endo, T.A., Okamoto, M., Nambara, E., Nakajima, M., Kawashima, M. et al. (2008) Arabidopsis Transcriptome Analysis under Drought, Cold, High-Salinity and ABA Treatment Conditions using a Tiling Array. Plant and Cell Physiology, 49, 1135–1149.

43 AlShareef, S., Ling, Y., Butt, H., Mariappan, K.G., Benhamed, M. and Mahfouz, M.M. (2017) Herboxidiene triggers splicing repression and abiotic stress responses in plants. BMC Genomics, 18, 260.

44 Liu, W., Duttke, S.H., Hetzel, J., Groth, M., Feng, S., Gallego-Bartolome, J., Zhong, Z., Kuo, H.Y., Wang, Z., Zhai, J. et al. (2018) RNA-directed DNA methylation involves co-transcriptional small-RNA-guided slicing of polymerase V transcripts in Arabidopsis. Nature plants, 4, 181–188.

45 Majewski, J. and Ott, J. (2002) Distribution and characterization of regulatory elements in the human genome. Genome research, 12, 1827–1836.

46 Rose, A.B., Elfersi, T., Parra, G. and Korf, I. (2008) Promoter-Proximal Introns in Arabidopsis thaliana Are Enriched in Dispersed Signals that Elevate Gene Expression. The Plant Cell, 20, 543–551.

47 Xu, Z., Wei, W., Gagneur, J., Clauder-Münster, S., Smolik, M., Huber, W. and Steinmetz, L.M. (2011) Antisense expression increases gene expression variability and locus interdependency. Molecular Systems Biology, 7, 468.

48 Andersson, R., Enroth, S., Rada-Iglesias, A., Wadelius, C. and Komorowski, J. (2009) Nucleosomes are well positioned in exons and carry characteristic histone modifications. Genome Research, 19, 1732–1741.

49 Chodavarapu, R.K., Feng, S., Bernatavichute, Y.V., Chen, P.-Y., Stroud, H., Yu, Y., Hetzel, J.A., Kuo, F., Kim, J., Cokus, S.J. et al. (2010) Relationship between nucleosome positioning and DNA methylation. Nature, 466, 388.

50 Nogués, G., Muñoz, M.J. and Kornblihtt, A.R. (2003) Influence of Polymerase II Processivity on Alternative Splicing Depends on Splice Site Strength. Journal of Biological Chemistry, 278, 52166–52171.

51 Vos, S.M., Farnung, L., Urlaub, H. and Cramer, P. (2018) Structure of paused transcription complex Pol II–DSIF–NELF. Nature, 560, 601–606.

52 Kumar, S.V. and Wigge, P.A. (2010) H2A.Z-Containing Nucleosomes Mediate the Thermosensory Response in Arabidopsis. Cell, 140, 136–147.

53 Birse, C.E., Minvielle-Sebastia, L., Lee, B.A., Keller, W. and Proudfoot, N.J. (1998) Coupling Termination of Transcription to Messenger RNA Maturation in Yeast. Science, 280, 298–301.

54 Prescott, E.M. and Proudfoot, N.J. (2002) Transcriptional collision between convergent genes in budding yeast. Proceedings of the National Academy of Sciences, 99, 8796–8801.

55 Hobson, D.J., Wei, W., Steinmetz, L.M. and Svejstrup, J.Q. (2012) RNA polymerase II collision interrupts convergent transcription. Molecular cell, 48, 365–374.

56 Fang, X., Wang, L., Ishikawa, R., Li, Y., Fiedler, M., Liu, F., Calder, G., Rowan, B., Weigel, D., Li, P. et al. (2019) Arabidopsis FLL2 promotes liquid–liquid phase separation of polyadenylation complexes. Nature.

