## Supplementary Figures for "Native Elongation Transcript sequencing reveals temperature dependent dynamics of nascent RNAPII transcription in *Arabidopsis*"

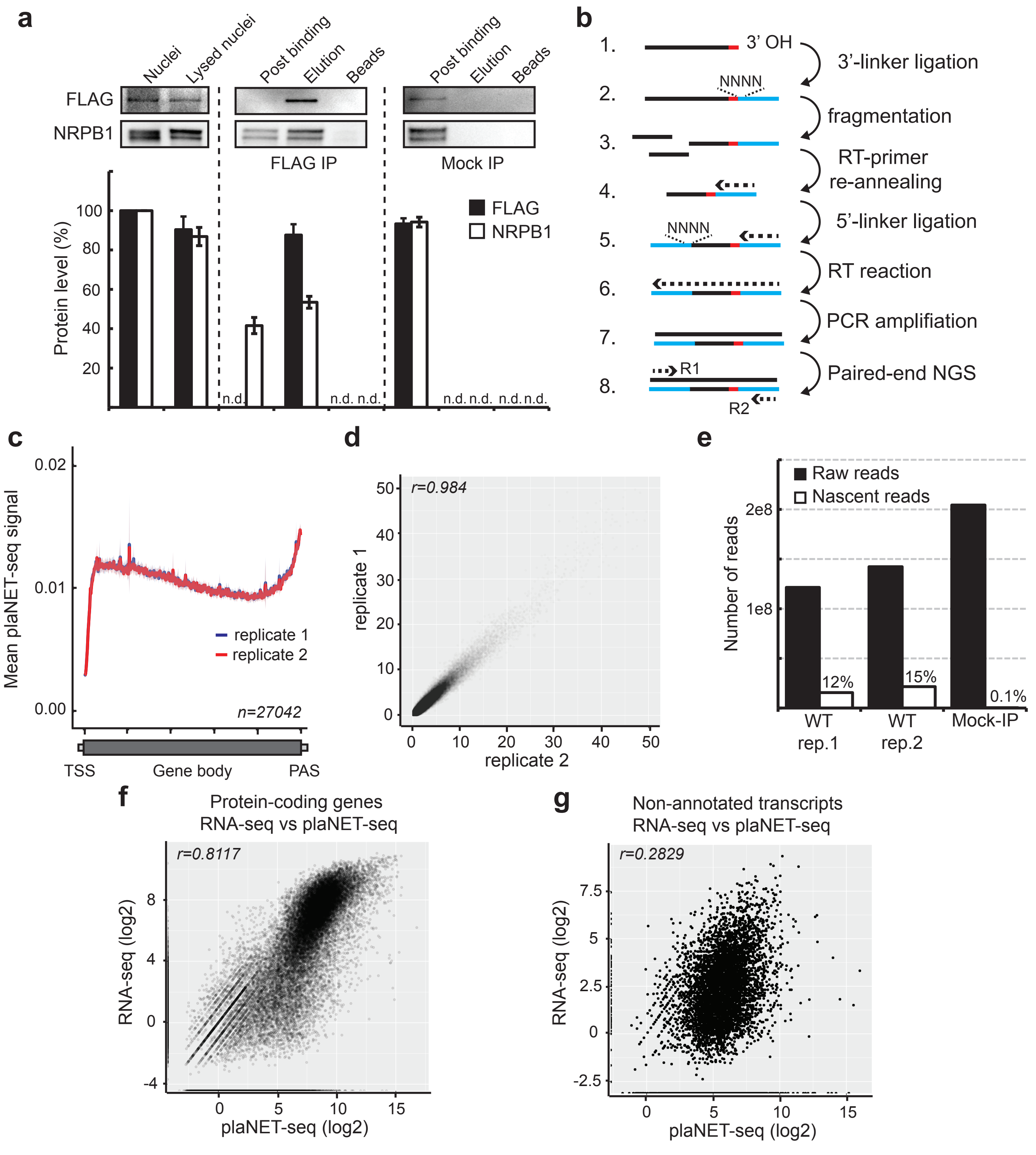


**Supplementary Figure 1:**
**a,** Western blots of NRPB2-FLAG and NRPB1 during the immunoprecipitation and elution steps of the plaNET-seq protocol. Upper panel shows representative blots (repeated with at least 3 biological replicates). Lower panel shows quantifications of proteins levels from blots. n.d. denotes non-detectable levels. Purification and elution of RNAPII complexes were efficient. A mock-IP showed no FLAG or NRPB1 signal in the elution, indicating high stringency of the purification.
**b,** Flow chart of the construction of plaNET-seq libraries. A 3'-linker was first ligated to the 3'OH (last base added by RNAPII, red dot) of nascent RNAs followed by alkaline fragmentation. Subsequently, a 5'-linker was ligated followed by an RT-reaction. A PCR reaction with barcode primers was performed before deep-sequencing. Both the 3'- and 5'-adapter contained 4 randomized bases that decreased the sequence bias of the RNA ligase and allowed for removal of PCR duplicates.
**c,** Metagene analysis of the plaNET-Seq signal over whole genes with length 0.5-5 Kb. Two biological replicates of untreated wild type sample are represented by blue (replicate 1) and red (replicate 2). The two replicates show very high reproducibility of metagene profiles. The shaded area shows 95% confidence interval for the mean.
**d,** Scatter plot of the reproducibility of plaNET-seq libraries. Correlation coefficient was determined using the Pearson method.
e, Bar chart showing raw and nascent reads (raw reads without PCR duplicates and contaminant reads) in two WT replicates and a mock-IP sample. The numbers denote the percentage of nascent reads compared to raw reads for each library. The mock-IP sample showed extremely low level of nascent reads (0.1% compared to 12-15% for WT replicates).
**f,** Scatter plot of the expression level of protein-coding transcripts as determined by strand-specific RNA-seq and plaNET-seq. Correlation coefficient was determined using the Spearman method. The correlation was high for protein-coding transcripts (i.e. mRNA).
**g,** Scatter plot of the expression level of non-annotated transcripts comparing strand-specific RNA-seq and plaNET-seq. Correlation coefficient was determined using the Spearman method. The correlation was low for non-annotated transcripts (i.e. lncRNA).

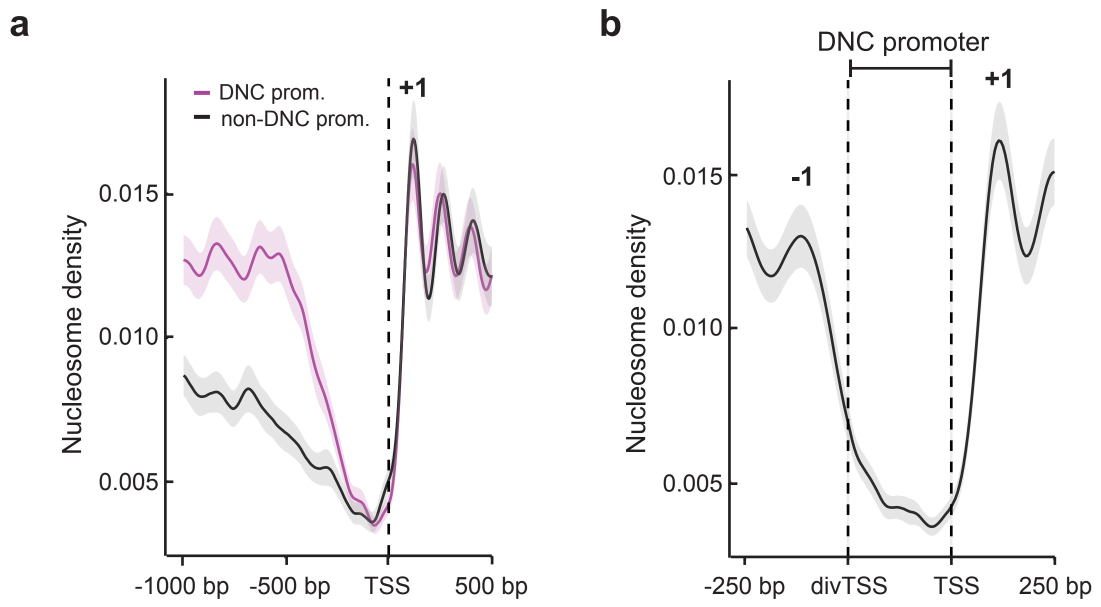


**Supplementary Figure 2:**
**a,** Metagene analysis of nucleosome density (DNC promoters versus non-DNC control promoters) in 1.5 Kb windows anchored at the sense TSS. The control promoters were chosen to match the DNC promoters by transcription level on the sense strand. The shaded area shows 95% confidence interval for the mean.
**b,** Metagene analysis of nucleosome density (DNC promoters only) in variable width windows anchored at both the sense TSS and at the divTSS (with 250 bp fixed width flanks). The shaded area shows 95% confidence interval for the mean.

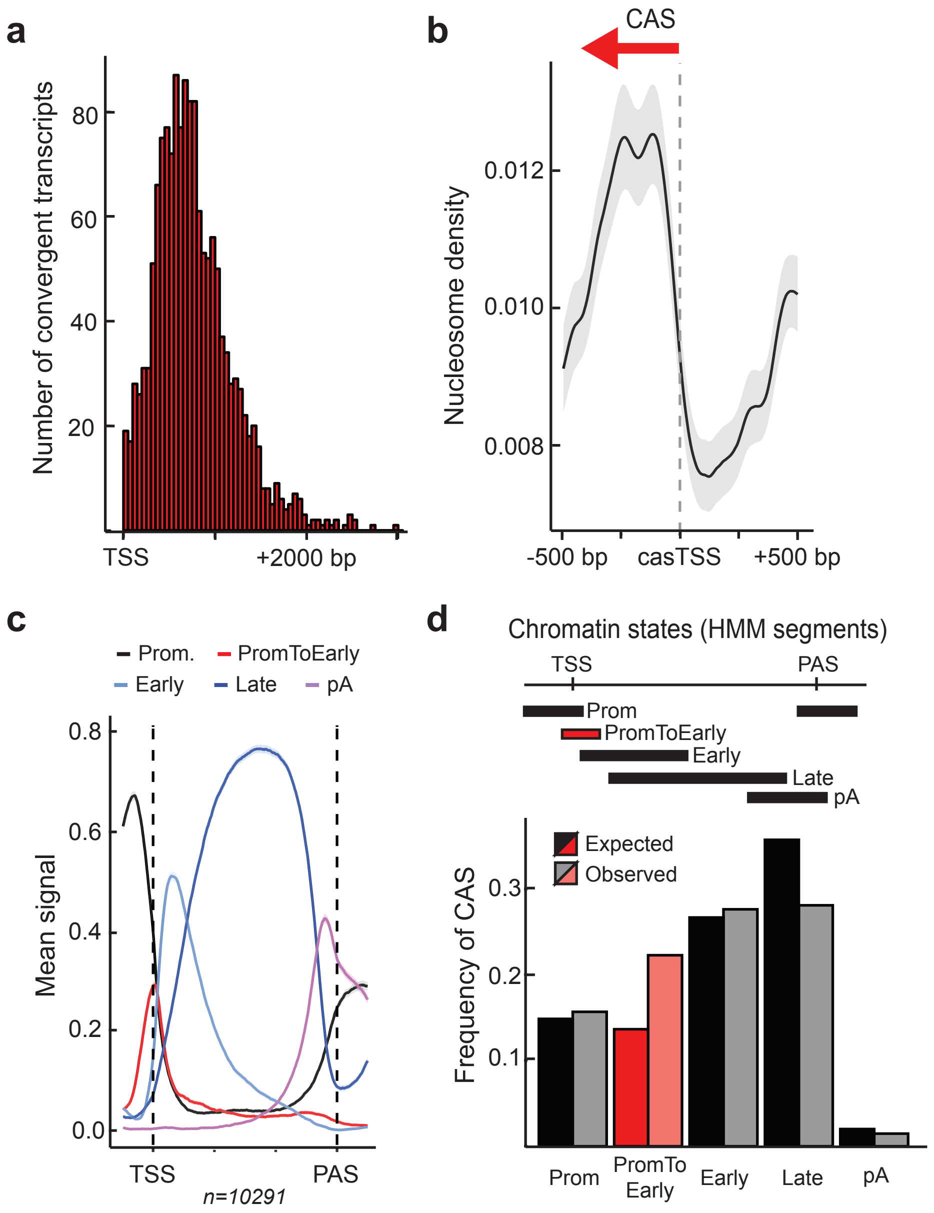


**Supplementary Figure 3:**

**a,** Absolute distance (bp) between the start site of convergent antisense transcripts (casTSS) and the sense TSS. CAS transcription tended to initiate within the first 1000 bp from the sense TSS.

**b,** Metagene analysis of nucleosome density in 1 kb windows centered at the convergent transcript start site (casTSS). The shaded area shows 95% confidence interval for the mean.

**c,** The distribution of chromatin state groups along nuclear protein coding genes (FPKM ≥ 1, length 1-5 Kb). The metagene plot covers the gene body (scaled to 300 bins) and includes 250 bp flanks upstream and downstream of TSS and PAS, respectively. The following groups of chromatin states from the PCSD database were defined: Promoter (Prom; states 13, 15-21), promoter-to-early elongation (PromToEarly; states 22-23), early elongation (Early; states 24-26), late elongation (Late; states 3-12, 27-28) and termination (pA; states 1-2).

**d,** Metagene analysis of chromatin states determined by ChromHMM along the gene bodies of Arabidopsis genes. Based on the PCSD database, the following states were assigned to respective group: promoter (Prom; states 13, 15-21), promoter-to-early elongation (PromToEarly; states 22-23), early elongation (Early; states 24-26), late elongation (Late; states 3-12, 27-28) and polyA (pA; states 1-2). Each CAS was assigned a chromatin state group based on overlap with casTSS. Observed frequencies of casTSS were plotted together with the expected frequencies of overlap based on the random model.

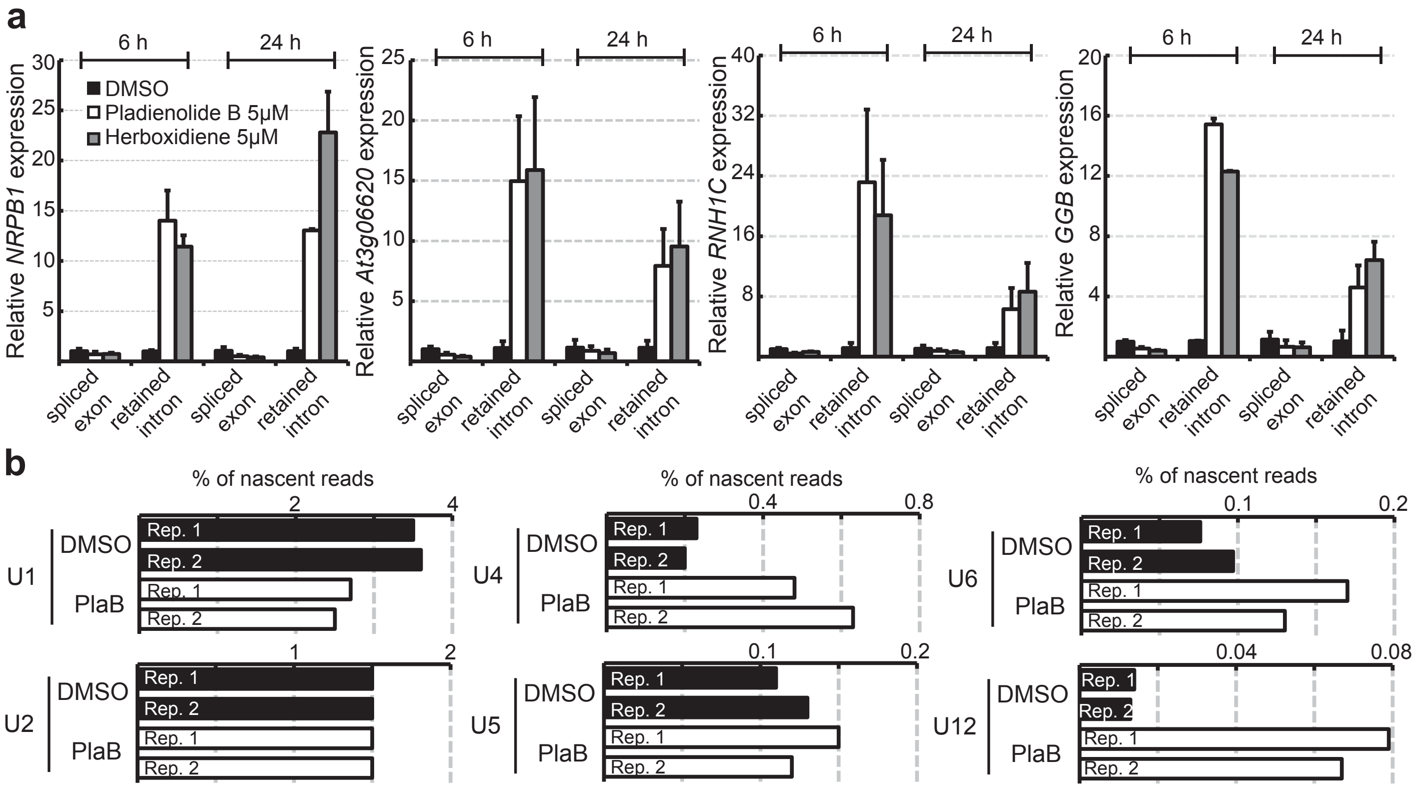


**Supplementary Figure 4:**
**a,** RT-qPCR confirmation of the plaB and Herboxidiene treatment (6h and 24h) shown for different splicing events (At4g35800, At3g06620, At1g24090 and At2g39550). Seedlings treated with DMSO for the same time were used as control. Bars represent mean ± SEM of three biological replicates.
**b,** Bar chart of the percentage of processed reads that aligned to different families of small nuclear RNAs involved in splicing. plaNET-seq DMSO and plaB replicates are shown.

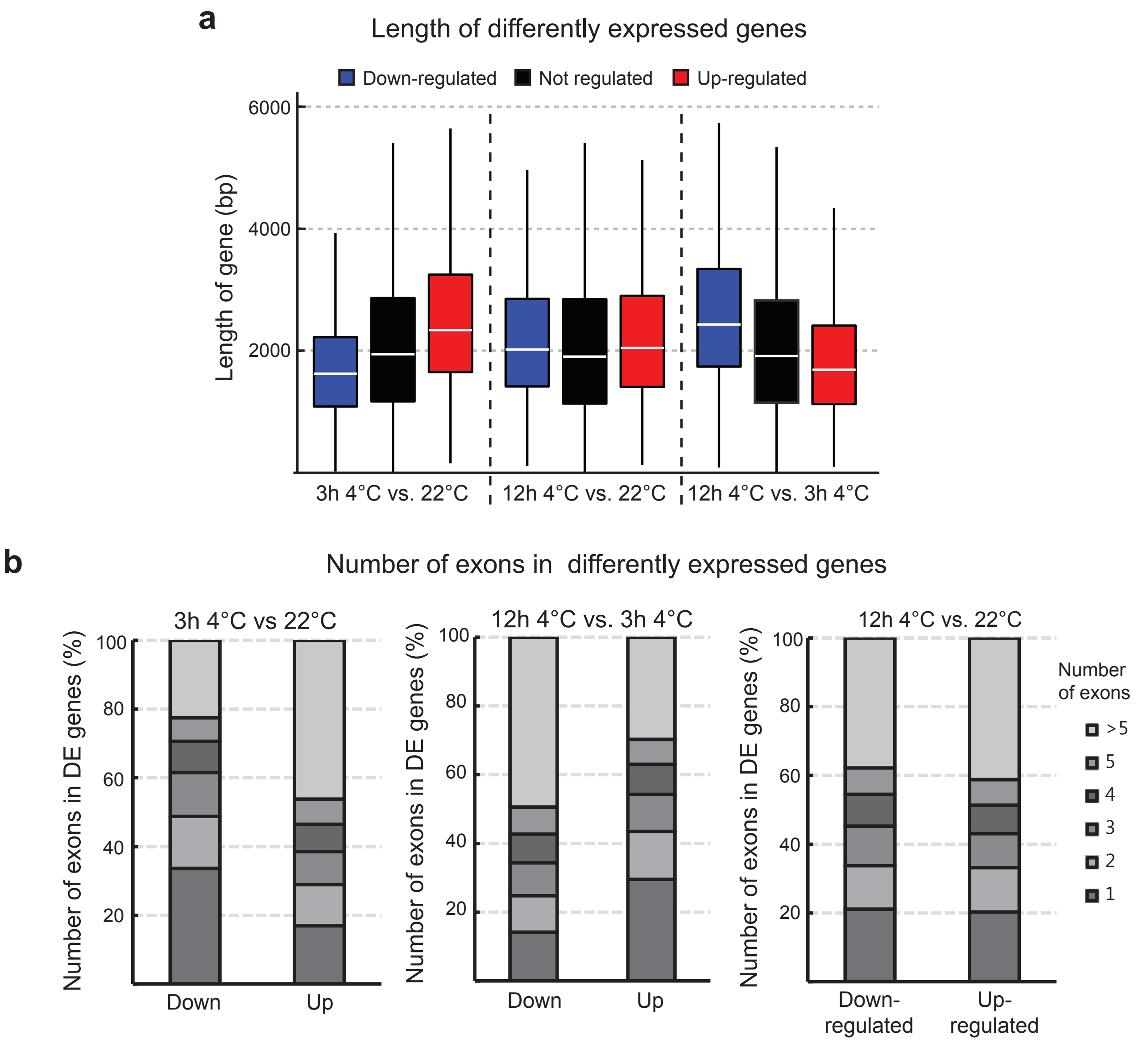


**Supplementary Figure 5:**
**a,** Box plots of the gene length in bp grouped by their differentially transcribed status in response to low temperature (as determined by plaNET-Seq). The plot shows genes which are differentially transcribed between 3h 4°C and 22°C (left), 12h 4°C and 22°C (middle) and 12h 4°C and 3h 4°C (right). Down-regulated genes are shown in blue, non-regulated in black and up-regulated genes in red. The graph shows that genes down-regulated after 3h at 4$^{o}$C tend to be shorter while up-regulated genes are longer. The opposite trend was detected between 12h at 4°C and 3h at 4°C. Here, down-regulated genes tend to be long while up-regulated genes are short.
**b,** Bar chart of the number of exons in genes differentially transcribed between 12h 4°C and 0h 4°C. Both down- and up-regulated genes show similar number of exons.

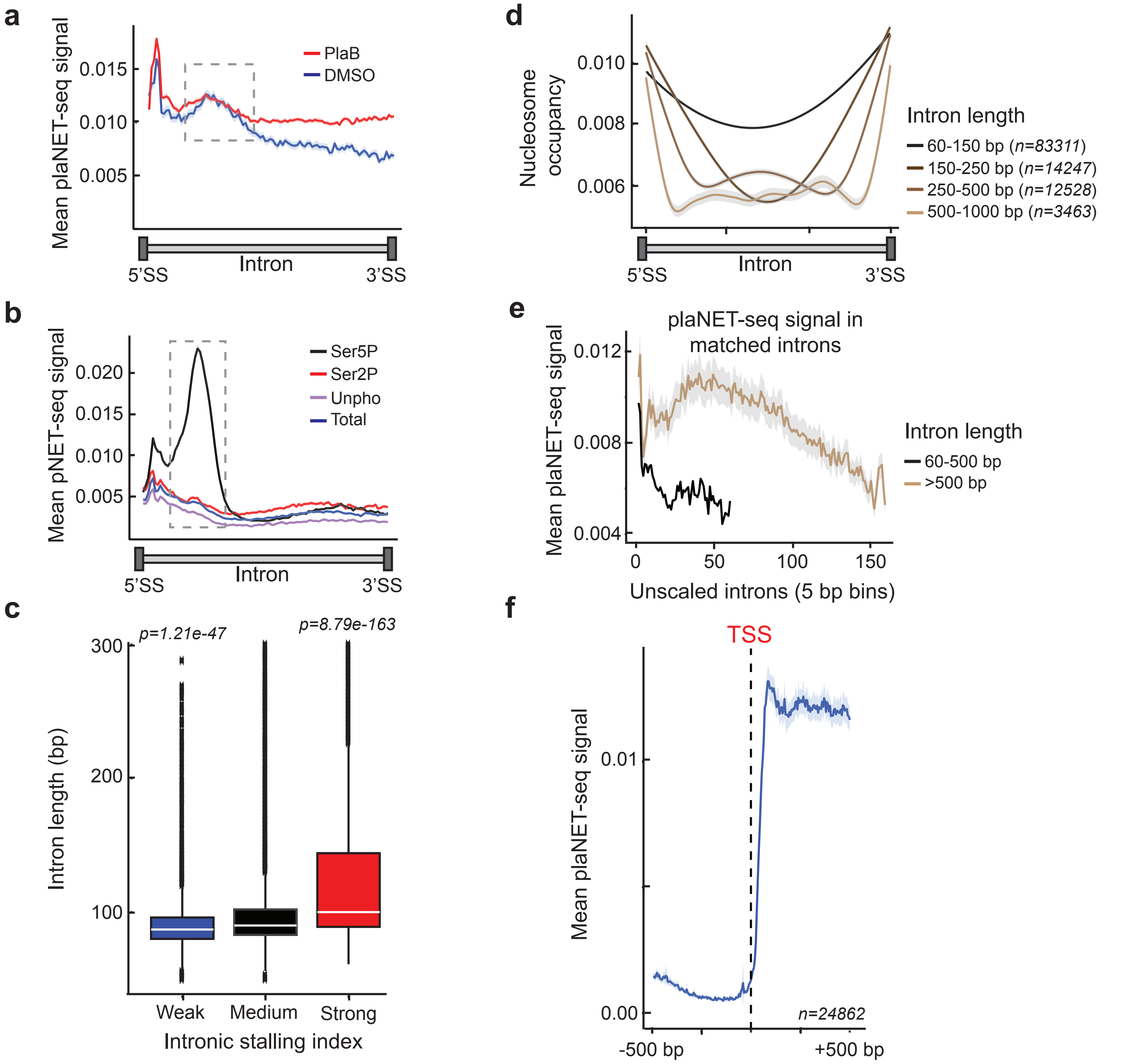


**Supplementary Figure 6:**

**a,** Metagene analysis of nascent RNAPII transcription in introns (with length 50-300 bp) as determined by plaNET-seq. DMSO is shown in blue and plaB in red. Dashed box indicates stalling site at the 5'-end of introns. Introns were scaled to 300 bins. The shaded area shows 95% confidence interval for the mean.

**b,** Metagene analysis of nascent RNAPII transcription in introns (with length 50-300 bp) as determined by pNET-seq. Data from the Ser5P antibody is shown in black, Ser2P in red, Unphosphorylated in purple and total RNAPII in blue. Dashed box indicates stalling site at the 3'-end of exons. The shaded area shows 95% confidence interval for the mean.

**c,** Box plots of intron length grouped by their intronic stalling index (ISI). Statistical significance of differences was measured by two-sided Mann-Whitney U test. Introns with strong intronic stalling tended to be longer and introns with weak stalling shorter compared to introns with medium stalling.

**d,** Metagene analysis of nucleosome occupancy in introns stratified by their lengths. Introns were scaled to 300 bins. Short introns (60-250 bp) were generally nucleosome-free while longer introns included one or more weakly positioned nucleosomes. The shaded area shows 95% confidence interval for the mean.

**e,** Metagene analysis of the plaNET-seq signal in matched introns stratified by length. Introns in both groups were chosen as pairs from the same genes to avoid any difference of transcription level. Longer introns showed a higher nascent RNAPII transcription, indicating a slower elongation compared to shorter introns. The shaded area shows 95% confidence interval for the mean.

**f,** Metagene analysis of the plaNET-seq signal in 1 b windows anchored at the annotated Transcription start sites (TSS) in wild type sample. No clear peak of RNAPII activity could be detected when the signal was anchored at the TSS. The shaded area shows 95% confidence interval for the mean.
